# HIF-dependent CKB expression promotes breast cancer metastasis, whereas cyclocreatine therapy impairs cellular invasion and improves chemotherapy efficacy

**DOI:** 10.1101/2021.09.09.459646

**Authors:** Raisa I. Krutilina, Hilaire Playa, Danielle L. Brooks, Luciana P. Schwab, Deanna N. Parke, Damilola Oluwalana, Douglas R. Layman, Meiyun Fan, Daniel L. Johnson, Junming Yue, Heather Smallwood, Tiffany N. Seagroves

## Abstract

The oxygen-responsive Hypoxia Inducible Factor (HIF)-1 promotes several steps of the metastatic cascade. A hypoxic gene signature is enriched in triple negative breast cancers (TNBCs) and correlates with poor patient survival. Since inhibiting the HIF transcription factors with small molecules is challenging, we sought to identify genes downstream of HIF-1 that could be targeted to block invasion and metastasis. Creatine kinase brain isoform (CKB) was identified as a highly differentially expressed gene in a screen of HIF-1 wild type and knockout mammary tumor cells derived from a transgenic model of metastatic breast cancer. CKB is a cytosolic enzyme that reversibly catalyzes the phosphorylation of creatine, generating phosphocreatine (PCr) in the forward reaction, and regenerating ATP in the reverse reaction. Creatine kinase activity is inhibited by the creatine analog cyclocreatine (cCr). Loss and gain of function genetic approaches were used in combination with cCr therapy to define the contribution of CKB expression or creatine kinase activity to cell proliferation, migration, invasion, and metastasis in ER-negative breast cancers. CKB was necessary for cell invasion *in vitro* and strongly promoted tumor growth and metastasis *in vivo*. Similarly, cyclocreatine therapy repressed cell migration, cell invasion, formation of invadopodia, and lung metastasis. Moreover, in common TNBC cell line models, the addition of cCr to conventional cytotoxic chemotherapy agents was either additive or synergistic to repress tumor cell growth.

## Introduction

A major clinical challenge in breast cancer is to treat metastatic disease. The overall survival of patients with metastatic breast cancer (MBC) remains dismal. Up to 30% of all patients will die within five years, and ~6% of patients are initially diagnosed with stage IV disease [1]. Targeting dysregulated tumor cell metabolism is a promising avenue to address drug resistance and to prolong the survival of patients [2,3]. Several metabolic pathways that are altered in tumors, including glutamine metabolism, fatty acid metabolism, and aerobic glycolysis are linked to therapeutic resistance [3].

The hypoxic response and the oxygen-responsive Hypoxia Inducible Factor (HIF) transcription factors play essential roles in mediating these pathways in tumors. HIF-1 and HIF-2 regulate genes that fine-tune cellular metabolism and that control cell proliferation, survival, or apoptosis [4,5]. HIF-1 is directly implicated in chemoresistance through regulation of metabolic input [5–7]. Hypoxia in general, and the HIFs, specifically, promote breast cancer metastasis and therapeutic resistance [7–9]. Using a transgenic model of MBC (MMTV-PyMT), we have shown that HIF-1α is essential for tumor growth and lung metastases originating from the mammary gland [9].

Targeting the HIFs with small molecules remains challenging. Several HIF inhibitors have been described, but most do not discriminate between HIF-1 or HIF-2, and they indirectly impact HIFα stability/activity [10]. Furthermore, deletion of either HIFα subunit is deleterious to normal development [11], suggesting potential toxicity. Finally, differential, or even competing roles, for HIF-1α and HIF-2α in tumorigenesis are reported [12].

To address these obstacles, we screened for genes downstream of HIF-1 that were differentially expressed after genetic knockout (KO) of HIF-1α to identify targets potentially more amenable to therapeutic intervention. Genes implicated in cellular metabolism and/or invasion, or for which chemical inhibitors had been previously identified, were prioritized for biological validation. One gene meeting both criteria was creatine kinase, brain isoform (CKB).

CKB is a member of a family of four creatine kinase (CK) enzymes that reversibly transfer a high-energy phosphate group between ATP and creatine, generating phosphocreatine and ADP [13]. Phosphocreatine is an essential local energy reservoir that highly metabolic tissues exploit to rapidly re-generate ATP from ADP to maintain a high ratio of ATP/ADP and to prevent local acidification near cellular ATPases [14,15]. These functions are relevant to tumors, which rely on aerobic glycolysis to produce energy and exist in an acidic microenvironment [6]. All four CK isoforms (two cytosolic, two mitochondrial) and the creatine transporter (*SLC6A8*) are direct HIF-responsive genes [16]. Neither CKB nor the mitochondrial CKMT1 are essential for development since single or compound knockout mice are viable [17]. CKB is over-expressed in several solid tumors including breast, colorectal, and ovarian [18–20]. In proteomic screens of prostate, lung, and HER2+ breast cancers, CKB was elevated [21–24].

Targeting creatine kinases is underdeveloped as an anticancer strategy targeting cellular metabolism. In the 1990s, several creatine analogs, including cyclocreatine (1-carboxymethyl-2-iminoimidazolidine, cCr), were shown to inhibit tumor growth with tolerable side effects [15]. Cyclocreatine has the most similar substrate kinetics to creatine, but unlike creatine, which is actively transported into cells by SLC6A8, cCr passively enters cells. cCr therapy was reported to repress rat mammary adenocarcinoma growth [25]; however, the impact on metastasis was not tested. Cyclocreatine also inhibits tumor cell motility *in vitro* [26]. In a colon cancer model, CKB was shown to mediate the metastatic potential of cancer cells by acting as a secreted kinase to produce phosphocreatine, which accumulated in the stroma and was then re-imported into tumor cells to promote CK-dependent survival to facilitate liver colonization [27]. In pancreatic adenocarcinoma, CKB was identified as a mechanosensitive transcriptional target of yes-associated protein 1 (YAP). Stiff substrates increased CKB levels and ATP production to promote collective cell invasion and chemotaxis [28]. These pre-clinical studies build upon *in vitro* observations that suggested CKB localizes in a spatially restricted manner in migrating cells to supply local ATP necessary for actin reorganization [29,30].

We tested if the creatine phosphagen arm of metabolism may be a key player downstream of the HIFs to promote breast cancer metastasis to the lung, a predominant site of metastasis in women with triple-negative breast cancer (TNBC) [31]. Using *in vitro* and *in vivo* approaches, including genetic modification of CKB compared to cCr treatment, we conclude that breast tumor cell-intrinsic expression of CKB is required to mediate cellular metabolism and to promote metastasis in ER-negative breast cancer models. Moreover, addition of cCr to conventional cytotoxic agents represses tumor growth in an additive or synergistic manner.

## Materials and Methods

### PyMT mammary tumor epithelial cells (MTECs)

HIF-1 WT and KO MTECs were maintained as in [9]. All cells were cultured at normoxia (ambient air) or exposed to hypoxia (0.5% O_2_) in a multi-gas incubator (acutely, ≤ 6h, or chronically, 16-24h).

### RNA extraction, gene expression microarray analysis and qPCR

RNA preparation and array processing protocols are reviewed in the Supplementary Methods. Raw data were transformed to the log_2_-scale, and normalized log-scale intensity values analyzed for differential expression using Expander. All genes with a mean fold-differential ≥ 2.0 (*p* <0.05) are included in gene lists (**Supplementary Tables S1-S3**), and can be assessed in GEO (GSE183694). qPCR was performed on a Roche LC480 instrument using primer and probe sets designed by the Universal Probe Library (UPL) assay design center; all primers are listed in **Supplemental Table S4**.

### Immunofluorescence of cultured cells

Cultured cells were stained using standard indirect immunofluorescence staining procedures, reviewed in the Supplementary Methods. A list of all antibody reagents is included in **Supplementary Table S5**. Slide images were captured on a Nikon inverted ECLIPSE Ti2 microscope using NIS-Elements software and the enhanced intensity projection module compiled images from multiple z-stacks.

### Protein extraction and western blotting

Cell extraction and western blotting was performed as in [9]. Western blotting details are included in the Supplementary Methods.

### Creatine kinase enzymatic activity assay

Protein extracts were analyzed by a CK assay (cat.#: C712-39, Pointe Scientific, Canton, MI) and activity was normalized to total protein content.

### Patient database mining

Please refer to the Supplemental Methods.

### Chromatin immunoprecipitation (ChIP) assays

Refer to Supplemental Methods for identification of HREs, sample preparation and ChIP analysis methods. Primers were designed to putative independent HRE sites, as well as to non-HRE sequences in the promoter regions (**Supplementary Table S6**). DNA was sheared to 500bp and ChIP was performed using antibodies against HIF-1@ or anti-rabbit IgG control and raw data analyzed as in [68].

### Generation of PyMT *Ckb* knockdown cells

Stable shRNA *Ckb* knockdown is described in the Supplemental Methods using targeting sequences listed in **Supplementary Table S7**. Two constructs, sh59 and sh61, produced >70% knockdown (**Supplementary Fig. S2A-B**).

### PyMT cell proliferation assay by WST-1

Cell growth in response to gene KD or to cCr treatment was measured as described in the Supplemental Methods.

### Wound healing assays

Wound healing assays were performed using a tip scratch method or using the wound healing protocol in 96-well format for the IncuCyte S3 live cell imager as described in the Supplemental Methods.

### Transient ectopic expression of CKB in MDA-MB-231 cells

MDA-MB-231-NR (NucLight red) cells (provided by Sartorius; RRID:CVCL_DF48) were transfected with FuGene HD reagent (Promega, Madison, WI) with 3 @g of either pCMV-6-Entry vector (OriGene, cat# PS100001, Rockville, MD) or pCMV-6-Entry expressing the CKB TrueORF (cat#RC203669); cells were selected with neomycin (2 mg/mL).

### Invasion assays

Invasion assays were performed as in [9], with modifications as described in the Supplemental Methods.

### Intracellular ATP Assay

Intracellular ATP levels were compared as in [69], with modifications as described in the Supplemental Methods.

### Seahorse bioanalyzer assays

PyMT cells were seeded into Seahorse XFe-96 sensor plates (Agilent Technologies, Santa Clara, CA) to produce near confluence within 18h. The next day, the plate was transferred to the Department of Pediatrics Bioenergetics Core for profiling on the XFe96 Extracellular Flux Analyzer. Cells were switched to an XF base medium supplemented with L-glutamine (2 mM) for the XF Cell Glyco Stress test or with glucose (10 mM), L-glutamine (2 mM), and sodium pyruvate (1 mM) for the XF Cell Mito Stress test. Cells were equilibrated for 1h prior to analysis of extracellular acidification rate (ECAR) or the oxygen consumption rate (OCR). After each run, cell number was quantified using CyQUANT dye (Thermo Scientific). Metabolic rates were calculated using the Seahorse XF report generator, and data imported into Prism 9.0 for analysis.

### Chemotaxis assays

Chemotaxis assays were performed in the IncuCyte S3 imager as recommended by Sartorius; additional experimental details are provided in the Supplemental Methods. MDA-MB-231-NR cells transiently transfected with pCMV-6-Entry +/− mCKB were seeded at a density of 1,000 cells/well of a ClearView chemotaxis plate (cat. #4582; n=4-6 replicates/cell line), which allows cell tracking in real-time. Cells were exposed to a reservoir containing DMEM +10% FBS as the chemoattractant and the chemotaxis software tool used to quantify cell migration. Raw data were normalized for initial plating density prior to export to Prism 9.0.

### Cell cycle analysis

Cell cycle analysis was performed as previously described [70]. Additional details are provided in the Supplemental Methods.

### PyMT primary tumor generation

PyMT EV, HIF-1 KO, sh59 *Ckb* KD, and sh61 *Ckb* KD MTECs were dissociated into single cells, diluted 1:1 in growth factor reduced Matrigel (BD Biosciences):HBSS, and injected into the cleared inguinal mammary fat pads of recipients (50,000 cells/10 μl). Tumor volume was measured with digital calipers [9]. Primary tumors were resected (@~500 mm^3^) in a survival surgery under anesthesia and mice allowed to survive until moribund due to metastasis.

### Tissue immunohistochemistry and quantification

Immunostaining was performed similar to [9]; additional details regarding quantification are provided in the Supplemental Methods.

### Tail vein assays in PyMT mice

PyMT EV, sh59, and sh61 *Ckb* KD cells (1×10^6^) were injected via the tail vein, mice were sacrificed after 21 days, and lungs inflated with formalin. Lung sections representing every 100 μm were stained with H&E and metastasis scored [9]. To test cCr efficacy, PyMT EV cells were injected, and allowed to seed the lungs for 24h. cCr dosing regimen was based on [25]. Mice were treated with vehicle (0.9% saline, daily, IP) or cCr (1g/kg, daily, IP) beginning day 1 post-injection. In an additional cohort, treatment with cCr was delayed until 7 days post-injection (after micro-metastases had formed) and on day 21 all mice were euthanized.

### Invadopodia assays using human TNBC cells

Assays are described in the Supplemental Methods. Briefly, glass coverslips were coated with OregonGreen-gelatin (cat. #G13186, ThermoFisher) [48]. Cells (BT549, 12,000/well or MDA-MB-231, 15,000/well) were seeded onto coverslips in growth media containing batimistat (10 μM; cat. # SML0041, Sigma) and incubated overnight. The next day, the media was replaced with fresh growth media to allow invadopodia to form, or cCr (25 mM) was also added. Plates containing seeded coverslips were placed into the IncuCyte imager and green fluorescence and phase contrast data collected (n=16 images/coverslip/time point). Green fluorescent area was normalized to seeding density per image and graphed as a ratio of change over time. At the end of the imaging, as optimized per cell line, coverslips were fixed and immunostained prior to mounting directly onto glass slides and imaged on a Nikon ECLIPSE Ti2 microscope.

### Growth inhibition during chemotherapy treatment

Live-cell imaging to enumerate human TNBC cells treatment in response to cCr monotherapy, or in combination with paclitaxel or doxorubicin, as well as calculation of the drug combination index (CI) is described in the Supplemental Methods.

### Statistical analysis

Unless otherwise stated, all data were entered into Prism 9.0 (Graph Pad, San Diego, CA) and analyzed using one-way or two-way ANOVA, followed by multiple pairwise comparison tests (*t*-tests). If standard deviation was not similar between two groups, Welch’s correction was applied. Graphs show the mean ± SEM and significance was determined with a 95% confidence level; *p*-values are indicated by asterisks, \**p*<0.05, \*\**p*<0.01, \*\*\**p*<0.001, \*\*\*\**p*<0.0001.

## Results

### *Ckb* is a HIF-1α dependent target in breast cancer

Tumor cell-intrinsic HIF-1α is required for mammary tumor growth and lung metastasis in the MMTV-PyMT mouse model [9]. To identify genes downstream of HIF-1 that mediate metastasis and may be more amenable to therapeutic intervention, we performed microarray profiling using HIF-1 WT and KO PyMT cells cultured at normoxia or 6h of hypoxia. Several hundred differentially-expressed genes were identified for the following comparisons: 1) HIF-1 WT normoxia vs. HIF-1 WT hypoxia; 2) HIF-1 WT normoxia vs. HIF-1 KO normoxia; 3) HIF-1 WT hypoxia vs. HIF-1 KO hypoxia (refer to Supplementary Tables S1-S3, respectively). Several genes down-regulated in HIF-1 KO cells, including *Egln3* (Phd3), *aldolase C* (Aldoc), and *Pdk1*, are known HIF-1α targets.

The enzyme creatine kinase brain isoform (CKB) was one of the top HIF-dependent, but not hypoxia-dependent, differentially expressed genes. *Ckb* was down-regulated in normoxic HIF-1 KO cells by 13.4-fold (p=0.0093) (Table S2). Following acute hypoxia exposure, Ckb expression in HIF-1 WT cells was 16.42-fold higher than in KO cells (p=0.0002) (Table S3). However, *Ckb* was not differentially expressed in HIF-1 WT cells between normoxic and hypoxic exposure (Table S1). Based on CKB’s well-described role in regulating energy metabolism, and prior studies using creatine analogs to inhibit the creatine kinase pathway, including cyclocreatine (cCr) [32], we sought to determine if CKB plays a key role in mediating breast cancer metastasis. Microarray profiling was performed using HIF-1 WT and KO cells derived from the PyMT model, since 100% of mice will develop lung metastasis on the FVB/Nj strain [33]. Likewise, primary tumors regenerated from HIF-1 WT PyMT tumor cells develop lung metastasis with 100% penetrance [9].

To validate Ckb expression changes, an independent set of PyMT HIF-1 WT and KO cells was exposed to normoxia or hypoxia. A significant decrease in Ckb mRNA expression was observed in normoxic HIF-1 KO cells compared to WT cells at either 6h or 24h of normoxia by qPCR, with a mean 7.44-fold decrease. At hypoxia, there was a more modest decrease in *Ckb* expression in the HIF-1 KO cells of ~1.7-fold (**Figure 1A**). We next investigated whether CKB is a direct HIF target in PyMT cells and in MCF-7 cells, which express CKB in an estrogen-responsive manner [34]. Two hypoxia-response elements (HREs) were identified in the murine *Ckb* or human *CKB* promoters (**Supplementary Figure S1A**). Chromatin immunoprecipitation (ChIP) assays were performed using PyMT HIF-1 WT and KO cells or MCF-7 cells modified to express pLKO.1-puro (empty vector, EV) or shRNA to HIF1A [35]. HIF-1α was recruited to the −1326 and −1835 sites in PyMT WT cells (**Supplementary Figure S1B**), with an enrichment in HRE site occupancy of 11.5-fold or 4.6-fold, respectively. There was 1.8 fold enrichment of HIF-1α at the −258 HRE site in MCF-7 EV cells relative to shHIF1A cells. However, no enrichment at the −935 site was observed (**Supplementary Figure S1C**). Therefore, HIF-1 directly regulates *CKB* mRNA expression in breast cancer cells, as reported for the colon [16]. Knockdown (KD) of *HIF1A* in MCF-7 cells significantly down-regulated *CKB* mRNA levels (**Supplementary Figure S1D**).

**Figure 1.**
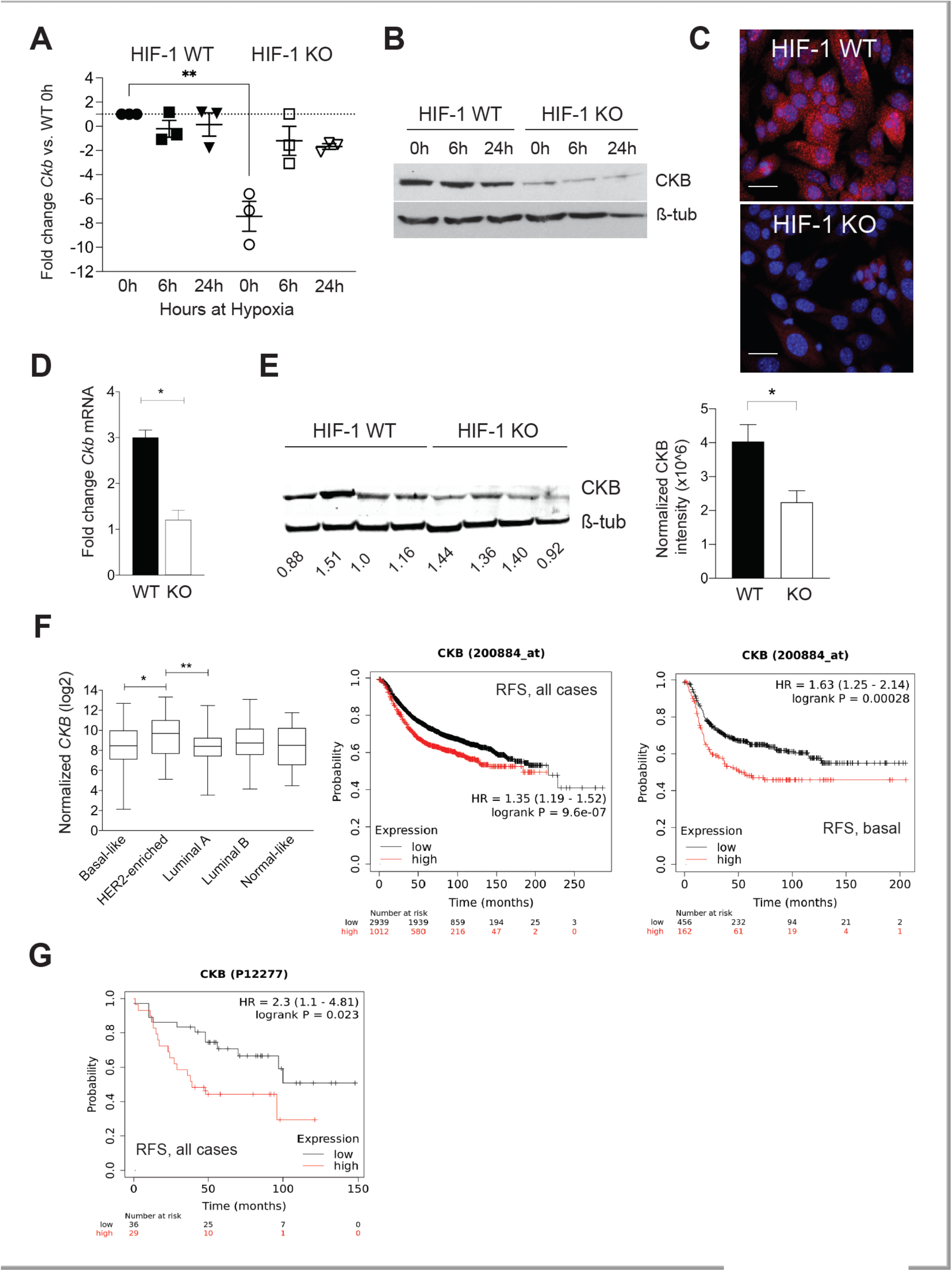
CKB expression in HIF-1 WT and KO PyMT cells and in end-stage tumors, and correlation to prognosis in clinical datasets. A. Scatter plot of the mean fold-change SEM in *Ckb* mRNA levels relative to HIF-1 WT cells at normoxia. Each data point represents an independent qPCR assay, and the grand mean is shown. B. PyMT cells were grown to 80% confluence and subjected to hypoxic culture for either 6h or 24h, or cells were continuously cultured at normoxia, such that the 0h sample was harvested on the same day as the 24h hypoxic sample. A representative western blot for CKB or ß-tubulin (loading control) is shown. C. Immunostaining for CKB (red) in HIF-1 WT and KO cells, counterstained with DAPI. Images were captured at 630x magnification (scale bar indicates 20 m). D. *Ckb* mRNA levels were compared in end-stage HIF-1 WT and KO tumors by qPCR. Data are expressed as the grand mean fold-change SEM between WT and KO tumors; the KO sample was set to a fold-change of 1.0 in each experiment. All data were first normalized for epithelial content based on *Krt18* (n=3 tumors/genotype/experiment). E. Western blotting for CKB comparing four independent tumors per HIF genotype (each harvested at similar volumes). The top half of the blot was blotted for CKB and the lower portion was blotted for ß-tubulin. Blots were imaged on a LiCor Odyssey system and CKB expression was compared in HIF-1 WT versus KO tumors after normalization to ß-tubulin. The bar graph shows the normalized CKB intensity SEM per genotype. F. (Left) TCGA was queried for *CKB* mRNA expression levels among breast cancer subtypes as defined by PAM50. Log_2_-normalized *CKB* expression is plotted, along with the 95% confidence interval and standard deviation. (Right) KM Plotter was used to plot the probability of RFS when patients were stratified by low or high expression of *CKB* mRNA levels (200884_at); plots are shown for either all breast cancer cases, or for basal breast cancer cases only. G. RFS is lower when CKB protein levels (P12277) are high. All images were directly exported from the KM Plotter tool.

CKB protein levels were also compared by western blotting. Exposure of HIF-1 WT PyMT cells to hypoxia for 24h did not induce CKB protein above normoxic (0h) levels (**Figure 1B**). Immunofluorescence also confirmed reduced CKB expression in HIF-1 KO PyMT cells (**Figure 1C**). CKB was localized to the cytoplasm in a perinuclear pattern, as well as to the nucleus. CKB association with the nuclear matrix has been reported in colon cancer cells [36]. CKB levels were then compared in PyMT HIF-1 WT and HIF-1 KO end-stage tumors [9]. A reduction in *Ckb* mRNA, normalized for epithelial content using keratin 18 (*Krt18*), (**Figure 1D**), and CKB protein in tumors was observed (**Figure 1E**). Of note, protein extracts were prepared from whole tumors, containing proteins expressed by tumor epithelial cells and the stroma, likely minimizing the total reduction in CKB since only the tumor epithelium lacks HIF-1α.

### CKB levels are correlated with shortened regression-free survival

The Cancer Genome Atlas (TCGA) database was queried to compare normalized *CKB* mRNA expression in breast cancer patients (**Figure 1F**). *CKB* is expressed in all subtypes, with enrichment in patients with HER2+ breast cancer relative to basal-like or luminal A subtypes. Comparing regression-free survival (RFS) using KMPlotter, high *CKB* mRNA expression correlated with a lower probability of RFS regardless of subtype (**Figure 1F**). In basal breast cancers, this hazard ratio increased (**Figure 1F**). Although limited breast cancer cases were available to query at the protein level, high levels of CKB also significantly correlated with reduced RFS (**Figure 1G**).

### Knockdown of *Ckb* impairs cell invasion

To determine if CKB impacts metastasis to impact overall survival, we generated *Ckb* loss of function models in PyMT cells using shRNA-mediated gene knockdown (KD). Two independent shRNAs produced >70% gene KD (herein referred to as “sh59” and “sh61”, **Supplementary Figure S2A**). Recombinant lentivirus particles produced for each shRNA construct were used to transduce HIF-1 WT cells, resulting in >70% reduction of *Ckb* mRNA and CKB protein compared to either HIF-1 WT or empty vector (HIF-1 WT+ EV) transduced cells (**Figure 2A-B** and **Supplementary Figure 2A-2B**). By qPCR and western blotting, the sh61 *Ckb* KD pool showed the greatest reduction (**Figure 2A-B**). Deletion of *Ckb* did not affect HIF-1α expression (**Supplementary Figure S2C**).

**Figure 2.**
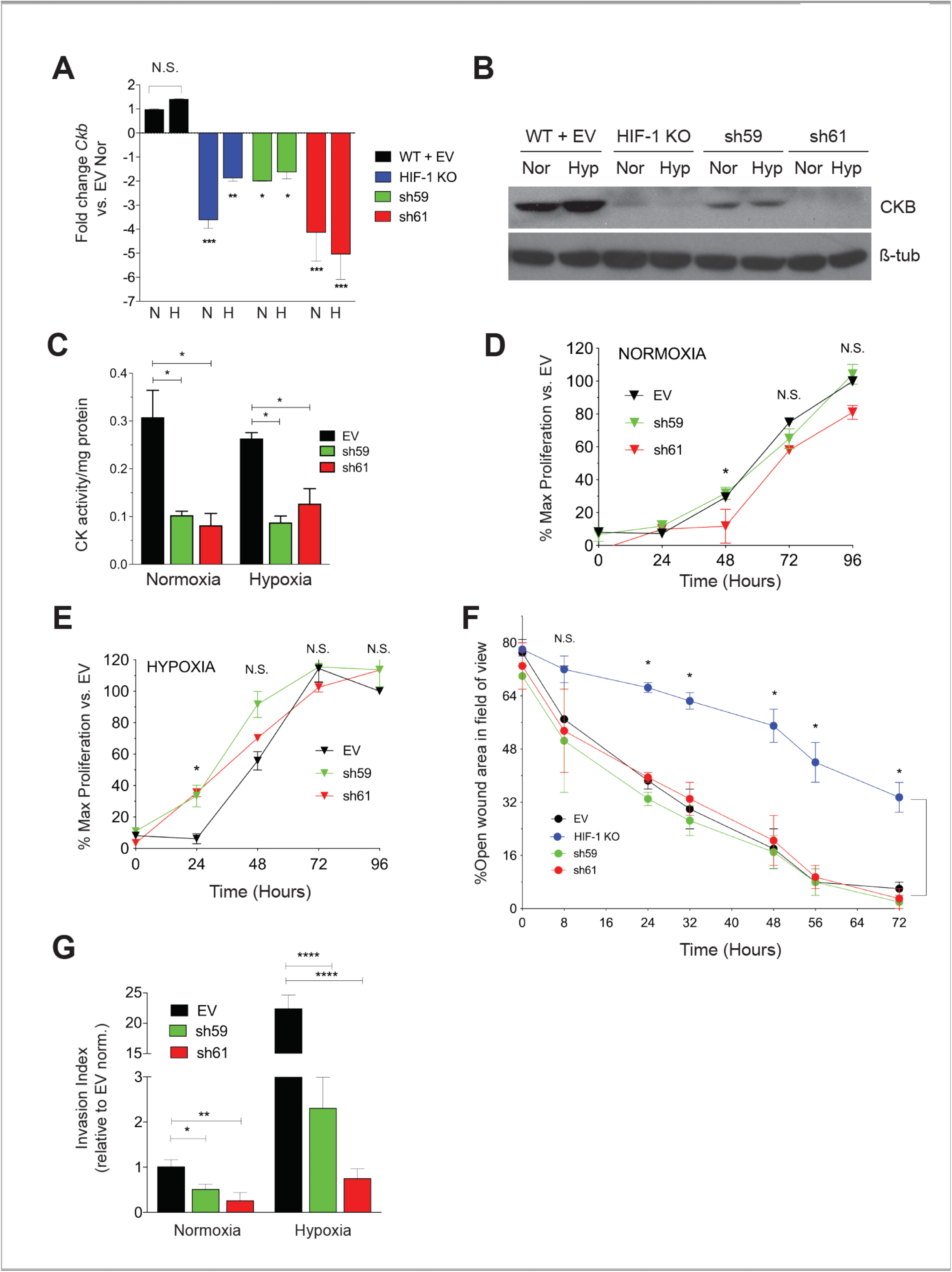
Effect of *Ckb* knockdown in PyMT cells upon cell growth and invasion. A. The mean fold change SEM in *Ckb* mRNA expression levels in HIF-1 KO cells and in shRNA KD pools (sh59 or sh61) relative to empty vector (WT +EV) cells as measured by qPCR. Cells expressing the sh61 shRNA showed the greatest reduction in *Ckb* mRNA levels. B. PyMT cells were grown to 80% confluence and subjected to normoxic or hypoxia culture for 6h and immunoblotted for CKB or ß-tubulin (loading control). Changes in CKB protein are consistent with changes observed by qPCR, with sh61 KD cells showing the greatest reduction of CKB expression. C. CK^act^ was measured in whole cell extracts prepared from PyMT EV, sh59, or sh61 cells cultured at normoxia or hypoxia (6h). Data shown are representative of three biological replicate experiments. D-E. Growth curves of PyMT EV, sh59, or sh61 cells cultured at normoxia (D) or hypoxia (E) in growth medium supplemented with 2% FBS. Cell proliferation was measured in replicate plates at each time point (24, 48, 72, or 96h) using the WST-1 assay. For both oxygen tensions, the grand mean of the percentage of proliferating cells relative to EV cells at t=0h SEM is presented, calculated as the average of mean cell number for n=3 technical replicates/time point/genotype as observed over three biological replicate experiments. All data were analyzed by two-way ANOVA; N.S.= not significant. F. PyMT EV, HIF-1 KO, sh59 KD, and sh61 KD cells were seeded into 6-well dishes such that they would be 100% confluent the next day. Following wounding, the percentage of open wound area in each field of view was measured and expressed as a ratio per the total area of the field of view. Data were analyzed by two-way ANOVA with a Bonferroni correction post-test. Data are representative of three biological replicate experiments. G. The grand mean fold change in the invasion index after data were normalized to EV cells cultured at normoxia (fold change set to 1.0; n=3 three independent biological replicate experiments). Data were analyzed by ANOVA with a Bonferroni post-test.

To determine if loss of CKB, only one isoform of the four creatine kinases in the cell, impacted total creatine kinase (CK) enzymatic activity, CK activity (CK^act^) levels were compared in extracts from PyMT EV and *Ckb* KD cells. Measuring CK^act^ is also important to predict sensitivity to cyclocreatine (cCr). Cells with very low CK^act^ (0.01-0.05 U/mg protein) are resistant or refractory to cCr, whereas cells with high CK activity are cCr-responsive (>0.10 U/mg protein) [37,38]. Independent of oxygen tension, loss of CKB reduced CK^act^ ~3-fold in sh59 and in sh61 KD cells (**Figure 2C**).

Cell proliferation was compared at either normoxia or hypoxia. At normoxia, there was a slight decrease in growth rate in sh61 KD cells, beginning at 48h (**Figure 2D**). However, at hypoxia, cell proliferation was significantly decreased in sh61 KD cells only at 24h (**Figure 2E**). There were no significant changes in proliferation observed between EV and sh59 KD cells. Overall, deletion of *Ckb* had minor effects on PyMT cell proliferation, similar to observations for PyMT HIF-1 KO cells [9].

We next compared breast cancer cell motility using the wound healing assay. There was no significant difference in wound closure between PyMT EV control cells and either sh59 or sh61 KD cells; in contrast, as expected, HIF-1 KO cells migrated significantly slower (**Figure 2F**). In contrast, invasion potential was significantly inhibited by *Ckb* KD at normoxia. Inhibition was more pronounced during hypoxia, with >10-20-fold reduction in invasion for shRNA KD cells relative to EV cells (**Figure 2G**).

### Loss of CKB represses glycolysis and oxidative phosphorylation

PyMT *Ckb* KD cells produced significantly less intracellular ATP than EV cells (**Figure 3A**). Metabolic activity was further characterized by Seahorse bioanalyzer assays. The extracellular acidification rate (ECAR), a measure of glycolytic activity, was reduced in either sh59 or sh61 KD cells (**Figure 3B-3D**). At the peak ECAR, glycolysis was repressed for each *Ckb* KD pool (**Figure 3C**), but only sh61 KD cells showed a reduction in glycolytic capacity (**Figure 3D**). Non-glycolytic acidification was also reduced in each KD pool (**Figure 3E**). These results are consistent with data from ovarian cancer cells that CKB is required for glycolysis [20]. In contrast, only the sh61 KD pool showed deficits in the oxygen consumption rate (OCR) (**Figures 3F-H**). Basal respiration levels and maximum respiration were reduced in the sh61 KD pool (**Fig. 3G-H**), although there was a non-significant trend in reduction for sh59 cells. ATP levels were significantly reduced in both shRNA KD pools (**Figure 3I**).

**Figure 3.**
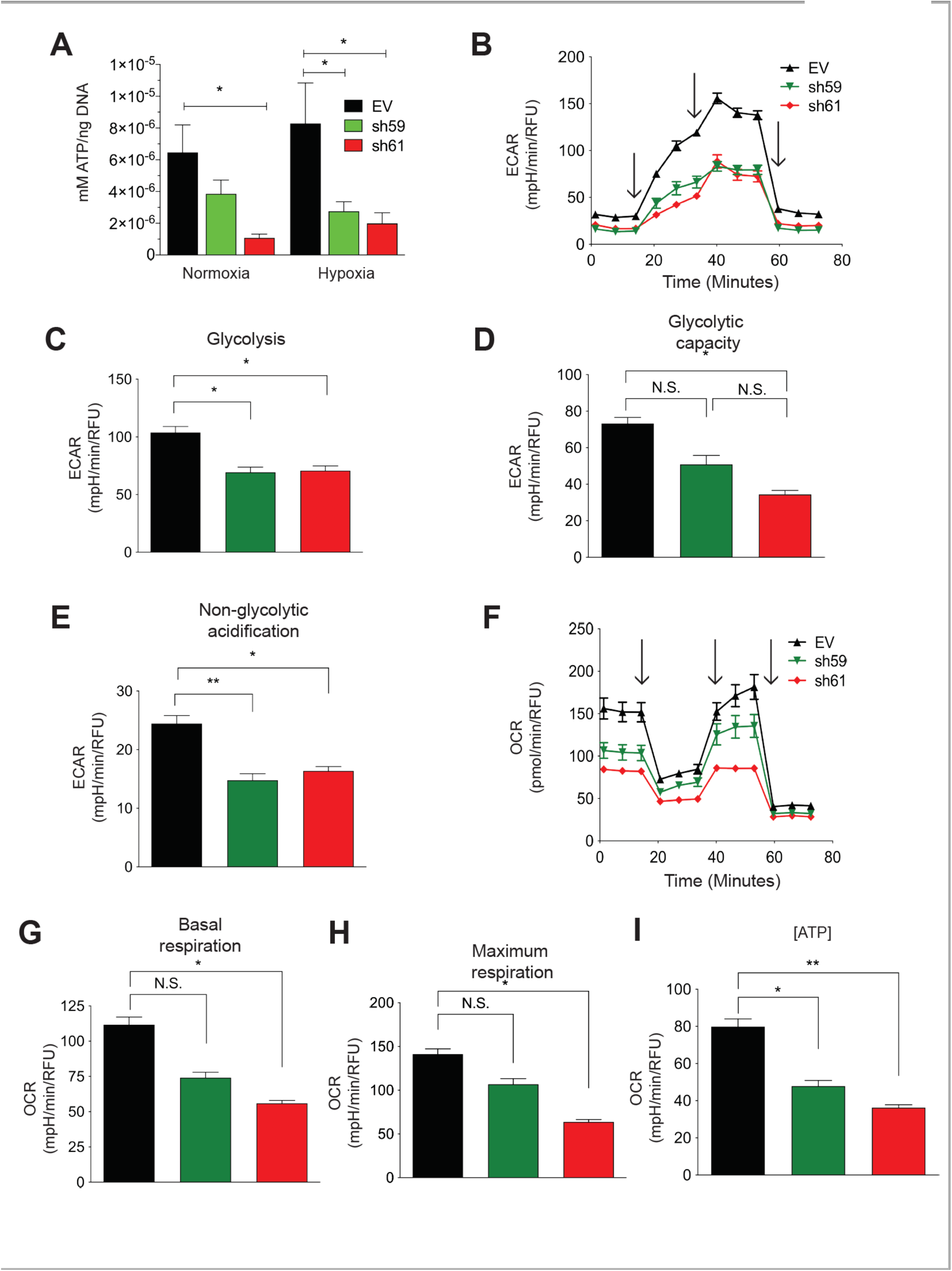
*Ckb* knockdown in PyMT cells reduces ATP levels and impairs both glycolysis and mitochondrial respiration. A. PyMT EV, sh59, and sh61 cells were plated in triplicate and grown to 80% confluence prior to normoxic or hypoxic culture (24h). Cells were harvested, washed with ice-cold PBS, and immediately lysed on the plate for comparison of intracellular ATP levels by a bioluminescent assay, and data normalized to total DNA content. The mean ATP concentration (mM) per ng of DNA SEM is shown (n=3 technical replicates per cell line/oxygen tension). The data shown are representative of three independent biological replicates. B-I. PyMT EV, sh59 *Ckb* KD, and sh61 *Ckb* KD cells were profiled for metabolic activity using Seahorse bioanalyzer assays (Glycolysis Stress or Mito Stress test kits) to measure changes in parameters associated with either ECAR, panels B-E, or with mitochondrial respiration (OCR, panels F-I). The black arrows (B, F) indicate when injections occurred during the assays. Changes were observed in ECAR plotted over time (B), peak glycolysis (C), glycolytic capacity (D), non-glycolytic acidification (E), OCR over time (F), basal respiration (G), maximum respiration (H), and ATP-linked respiration (I) in sh61 *Ckb* KD cells. For some of the measured outputs, there were also significant changes observed in sh59 *Ckb* KD cells. Panels B and F are representative of three independent experiments. For panels C-E and G-I, the bar graphs represent the grand mean of three independent experiment means across n=6 technical well replicates/genotype/experiment. Each genotype of cells was randomly plated in different patterns during each independent experiment to minimize any potential effects of plate well location on measurements.

### Ectopic CKB expression promotes, but cyclocreatine treatment represses, cell migration and invasion

Mouse CKB (mCKB) was stably transfected into HIF-1 KO PyMT cells to determine if re-expression of CKB would rescue invasion. The percentage of cells invading through ECM increased when CKB was re-expressed (**Figure 4A**). CKB was also ectopically expressed in a highly invasive TNBC model, MDA-MB-231 cells, which do not express detectable levels of CKB protein by immunostaining (data not shown) or by immunoblotting (**Figure 7A**). After transient transfection, the percentage of invading cells increased almost 3-fold (**Figure 4B**), with similar results when mCKB was stably expressed (data not shown). Next, MDA-MB-231-NucLight Red (NR) cells, in which nuclei are labeled with a dye to allow for real-time imaging during chemotaxis, were transiently transfected with vector or +mCKB. Cells were exposed to reduced serum overnight (2% FBS) and then seeded into ClearView plates and attracted towards 10% FBS. Over-expression of mCKB in MDA-MB-231-NR cells dramatically increased chemotactic potential (**Figure 4C**).

**Figure 4.**
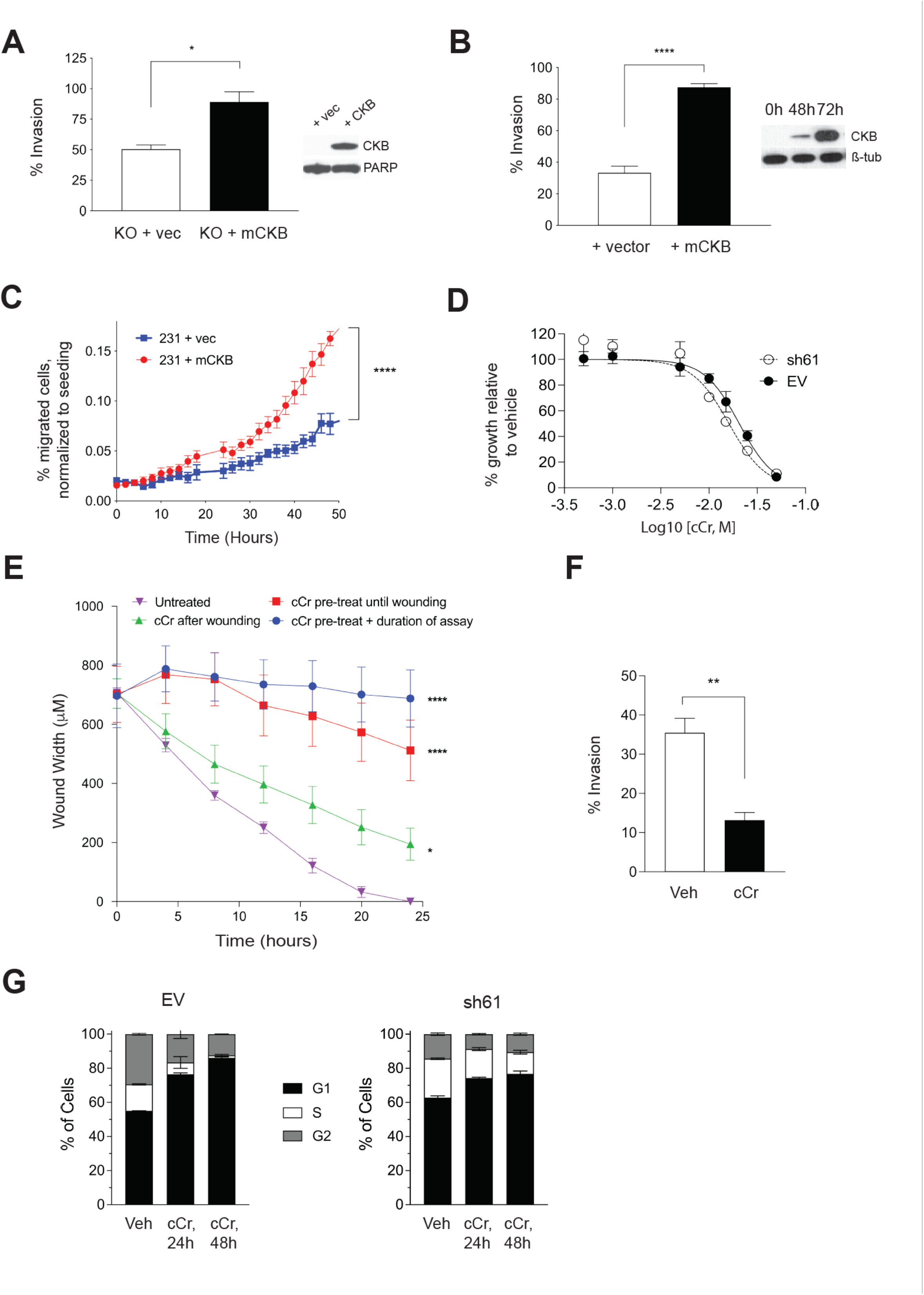
Ectopic expression of CKB in HIF-1 KO PyMT cells or in human MDA-MB-231 TNBC cells enhances cell invasion, but cCr treatment represses wound healing, invasion, and cell cycle progression. A. The percentage of invading cells was compared between PyMT HIF-1 KO + vector control (+ vec) and PyMT HIF-1 KO + mCKB (+CKB) cells over three biological experiments; the grand mean SEM is shown. The insert shows CKB expression by western blotting relative to total PARP (loading control). B. The percentage of invading cells in MDA-MB-231 TNBC cells transfected with vector alone (+ vector) or with mCKB (+ mCKB); cells were plated for invasion assays 72h post-transfection, when CKB levels were maximally expressed (western blot insert, ß-tubulin as a loading control). C. Chemotaxis assays were performed in transiently transfected MDA-MB-231-NR cells using IncuCyte ClearView plates. The mean percentage of migrated cells/total cells seeded was compared between genotypes over time during chemoattraction to 10% FBS. Data are representative of two independent experiments. D. Cells were exposed to increasing doses of cCr as described in the methods and imaged for 96h. Percent growth inhibition was set relative to vehicle control and then plotted vs. log of the molar concentration and fit with a nonlinear regression curve analysis. The graph is representative of at least 3 independent biological replicates. E. PyMT EV cells were plated onto ImageLock plates. Then, cells were either pre-treated with cCr prior to wounding and cCr added post-wounding, cells were pre-treated with cCr prior to wounding, but cCr removed post-wounding, or fresh media with cCr was added only after wounding. Data are representative of at least two biological replicates. F. PyMT EV cells were seeded in an invasion assay in which cells in the upper chamber were exposed to either vehicle or to 25 mM cCr and then cultured for 48 h. G. PyMT EV or sh61 cells were treated with 25 mM cCr for either 24h or 48h and cell cycle analysis was performed by PI staining to assay for changes in cell cycle progression. The mean SEM is shown for each phase of the cell cycle (n=3 technical replicates/genotype/time point). Data are representative of three independent experiments.

To test whether treatment with cyclocreatine (cCr) would mimic loss of CKB function, EV and sh61 *Ckb* KD PyMT cells were treated with increasing concentrations of cCr for 96h and growth inhibition evaluated. We had expected to observe that cCr sensitivity would be reduced in *Ckb* KD cells, but, dose response curves were overlapping. The IC_50_ value in EV and sh61 cells at 96h was 13.73 ± 4.33 mM and 16.43 ± 2.98 mM, respectively (**Table 1**). Since 15 mM and 25 mM cCr doses were then used in short term biological assays (24-48h duration), we confirmed that these doses do not induce extensive cell death in EV or HIF-1 KO cells compared to vehicle treatment, although the basal level of cell death is higher in HIF-1 KO cells (**Supplementary Figure S3A-B**).

**Table 1.** Cyclocreatine (cCr) IC_50_ values in PyMT ER-negative cells. IC_50_ values ± SEM were determined at 96h as described in the methods. IC_50_ values for independent experiments were calculated using non-linear regression and variable slope fit in Prism 9.0. Results from independent experiments were averaged to report the mean (n=6, WT; n=6, HIF-1 KO; n= 4, EV; n=6, sh61).

| HIF-1 WT | HIF-1 KO | EV | sh61 CKB KD |
| --- | --- | --- | --- |
| 16.76 mM ± 4.47 | 15.44 mM ± 3.95 | 13.73 mM ± 4.33 | 16.43 mM ± 2.98 |

We tested cCr for anti-metastatic activity in a high-throughput wound assay. Either pre-treatment of PyMT EV cells with cCr prior to seeding, or adding cCr only after the scratch was applied inhibits wound healing (**Figure 4E**). The most potent repression occurred when cells were pre-treated with cCr 24h prior to wounding and then cCr was re-added during healing. However, cCr pre-treatment before wounding without adding cCr post-scratch also significantly delayed wound healing (**Figure 4E**). Exposure of EV cells to cCr also represses invasion through Matrigel (**Figure 4F**). Overall, small molecule inhibition of the CK pathway via cCr mimics loss of CKB function.

The chemotherapeutic potential of cCr is also related to its ability to block cell cycle progression. Short-term treatment with cCr has reversible effects on the cell cycle, whereas long-term treatment leads to cell death [39]. EV cells treated with cCr accumulated in the G1 phase after 24h, with a corresponding reduction in the percentage of cells in the S-phase; these results were exacerbated at 48h **(Figure 4G**). In contrast, sh61 KD cells showed smaller increases in the G1 phase or decreases in the S-phase and the differences between 24h and 48h of cCr treatment were negligible (**Figure 4G**).

### Loss of *Ckb* suppresses tumor growth and inhibits lung metastases

We compared growth of mammary tumors regnerated from orthotopic implantation of PyMT EV, HIF-1 KO, sh59 KD, and sh61 KD cells. In contrast to in vitro cell proliferation results, tumor growth rate was suppressed and was very similar between HIF-1 KO tumors and each Ckb shRNA pool (**Figure 5A**). There was no significant difference in tumor volume between each shRNA pool. At day 34 post-transplant, the mean tumor volume of EV cells was ≥1000 mm^3^, whereas all other genotypes had reached a maximum volume of ~200 mm^3^. Decreases in tumor wet weight were similarly observed (**Figure 5B**). To investigate if loss of Ckb in the tumor epithelium impacted overall survival, in an independent experiment, all tumors were grown until they could be resected at a similar volume, and animals were housed until moribund due to lung metastases. Animals previously bearing CKB+ EV tumors became moribund >3x faster than recipients of shRNA KD cells (HR= 3.681, **Figure 5C**). Tumor sections were then immunostained with antibodies to either Ki67, activated caspase-3 or CD31 and area positive for these markers quantified. Likely because PyMT EV tumors are highly necrotic, the percent positive area for Ki67+ cells was higher in all of the smaller tumors ((HIF-1 KO, sh59, and sh61, **Figure 5D**). As expected, CD31 levels were decreased in HIF-1 KO tumors relative to EV tumors, whereas a reduction in CD31 was only significant for sh61 KD tumors (**Figure 5E**). Approximately 9.8% of EV tumor area was apoptotic. In contrast, very low levels of apoptosis were observed for HIF-1 KO tumors (0.77%), and apoptosis was lower in shRNA KD tumors compared to EV tumors (**Figure 5F**).

**Figure 5.**
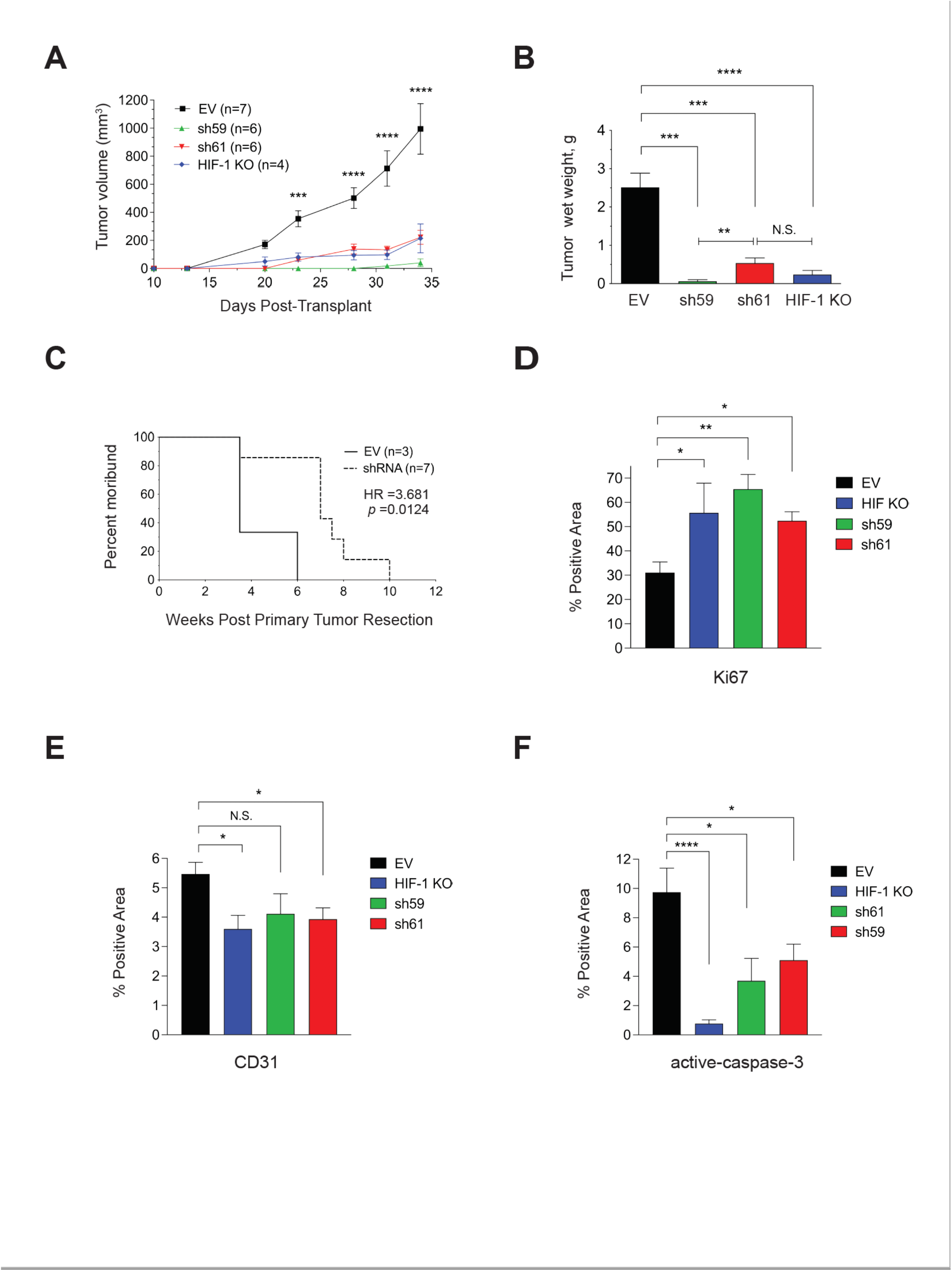
Tumor cell-intrinsic CKB promotes PyMT primary tumor growth and lung metastasis *in vivo* and deletion of CKB in the tumor epithelium improves overall survival. A. Growth rate over time after transplantation of PyMT EV, HIF-1 KO, sh59 KD, and sh61 KD cells into the inguinal mammary fat pad of female FVB/Nj recipients. Mean tumor volume ±SEM is shown. B. All tumors harvested at day 34 from panel A were weighed to determine the mean wet weight (g)± SEM at experiment endpoint. Data in A-B are representative of two independent experiments. C. The impact of CKB expression in the PyMT tumor epithelium on the survival of recipients following tumor resection. Mice implanted with PyMT cells (EV or either shRNA KD construct) were subjected to primary tumor resection after tumors grew to similar volumes (~500 mm^3^) and mice allowed to survive post-resection until moribund due to lung metastasis. Mice were removed from the study when panting due to lung metastasis, or if bodyweight decreased by >15%. The morbidity hazard ratio (HR) is 3.68 times higher when FVB/Nj recipients bear *Ckb* WT tumors (n=3 mice for EV and n=7 total mice for *Ckb* shRNA knockdown, representing either sh59 KD or sh61 KD tumors). D-F Immunostaining of PyMT EV (n=5 tumors), HIF-1 KO (n=4 tumors), sh59 KD (n=4 tumors) or sh61 KD (n=5 tumors) sections to enumerate Ki67 (D), CD31 (E) or activated-caspase 3 (F). The mean SEM of the percentage (%) of the positive area of whole tumor sections is reported for each genotype.

### Either *Ckb* deletion, or systemic treatment with cyclocreatine, blocks lung metastasis

To explore if *Ckb* knockdown blocks latter stages of the metastatic cascade, PyMT EV, sh59, or sh61 KD cells were injected into the tail vein to generate lung metastases. Lungs of mice injected with EV cells were filled with large metastases visible to the naked eye, whereas no surface metastases were observed for sh61 KD cells (**Figure 6A**). A strong repression of metastasis was observed for both shRNA *Ckb* KD pools, but there was no significant difference between the two pools (**Figure 6A**).

**Figure 6.**
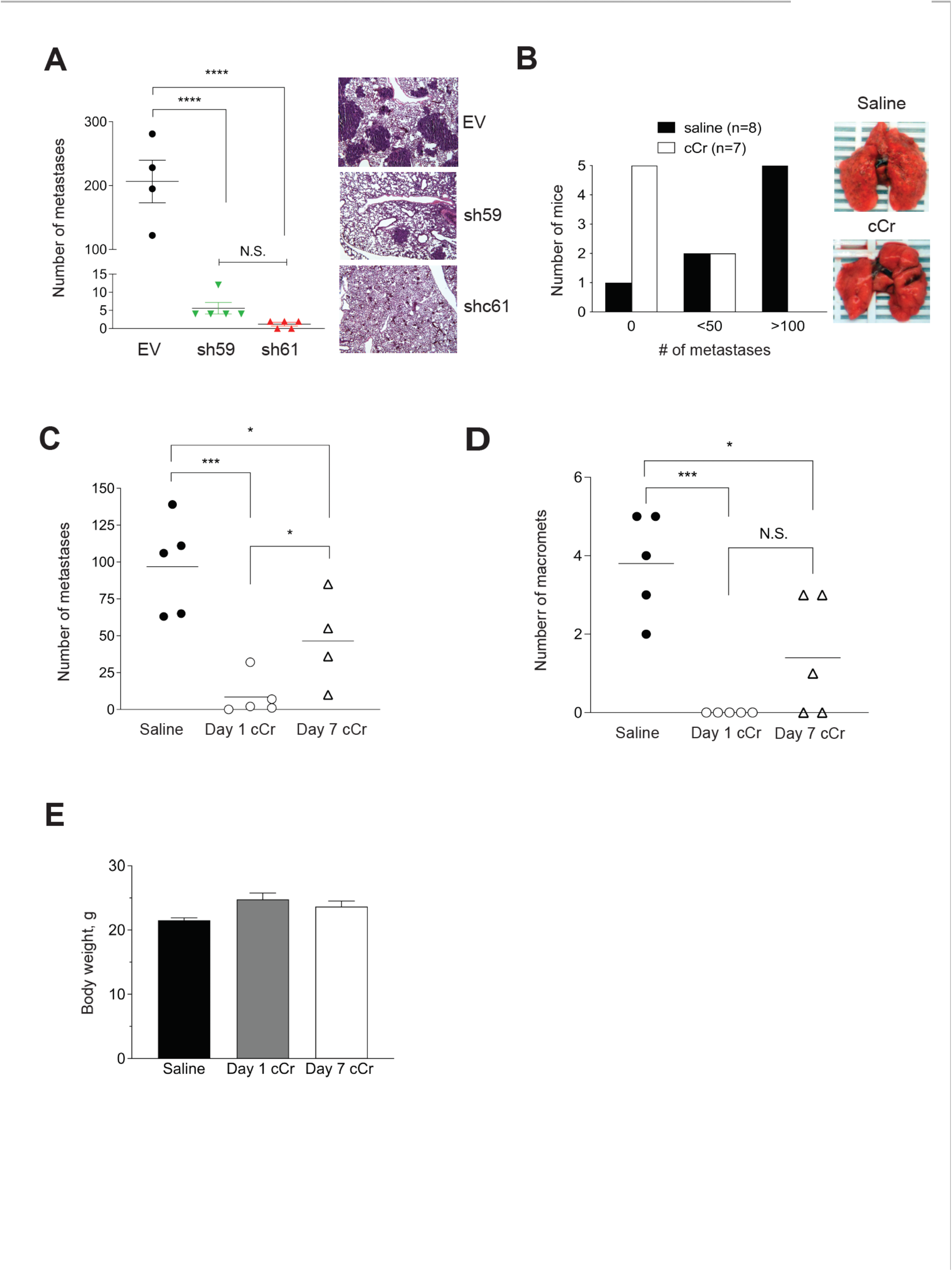
Knockdown of *Ckb* in the PyMT tumor epithelium or systemic cCr therapy decreases lung metastasis burden. A. PyMT EV, sh59 KD, or sh61 KD cells were injected into the tail vein of female FVB/Nj recipients. After 21 days, mice were euthanized and lungs harvested for metastasis evaluation after lung inflation through the trachea with PBS. The number of metastases was compared across genotypes; the scatter plot shows the burden of individual lungs with the mean ± SEM shown. Corresponding H&E--stained images of lungs representative of the genotype mean are shown (400x magnification). Data are representative of two independent experiments. B. EV PyMT cells were injected into the tail vein of female FVB/Nj recipients. The next day, treatment with either vehicle (saline, IP, daily) or cCr (1g/kg in saline, IP, daily) was initiated. After 21 days, the mice were euthanized and lungs harvested after inflation with PBS. The total number of surface metastases was counted under a dissecting scope. A majority of the mice treated with cCr (5 of 7) did not develop detectable metastases, whereas 2 mice in each cohort (2/8, saline and 2/7, cCr) developed fewer than 50 metastases. Only in the saline group did the majority of mice develop metastases throughout the lung, with >100 lesions present (5/8 mice; c2= 7.634, *p*=0.022). Images of whole lungs photographed immediately post-dissection are shown. C-D. Comparison of total metastases present per lung (C), or the number of macro-metastases per lung (D) when mice are treated with vehicle, or when cCr is administered at either day 1 (Day 1 cCr) or when cCr therapy begins at day 7 post-tail vein injection (Day 7 cCr). All lungs were harvested at day 21 post-injection. E. The mean bodyweight of the mice treated with saline (vehicle) or cCr from panels C-D.

**Figure 7.**
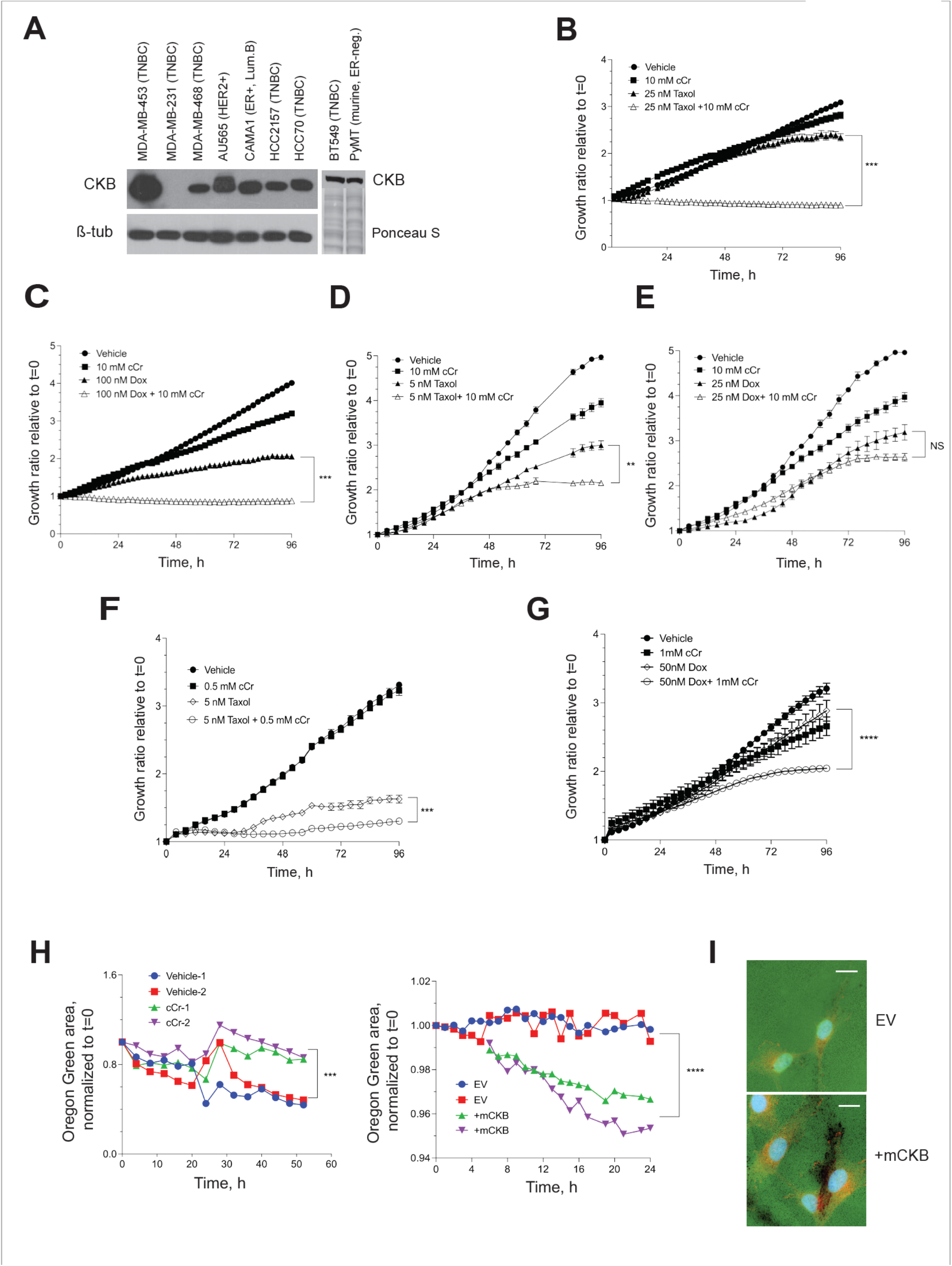
Cyclocreatine is synergistic with, or additive to, two conventional chemotherapies, and cyclocreatine inhibits the formation of invadopodia. A. (Left) Western blotting for endogenous CKB protein in a panel of human breast cancer cell lines (ß-tubulin, loading control). (Right) Western botting for CKB protein in BT549 TNBC cells and HIF-1 WT PyMT cells in a separate blot. BT549 cells express similar levels of CKB protein as PyMT HIF-1 WT cells; equivalent loading is indicated by Ponceau S staining. B-G. The growth ratio (all treatments are normalized to their respective t=0 cell density) over time when human MDA-MB-468 TNBC cells (B-C), BT549 TNBC cells (D-E), or MDA-MB-453 TNBC cells (F-G) are cultured in the presence of vehicle, cCr alone, or cCr with Taxol (B, D, F) or doxorubicin (C, E, G), for 96h. H. Loss of Oregon Green-conjugated gelatin was measured in the IncuCyte S3 live-cell imager after seeding cells onto gelatin-coated coverslips in the presence of an MMP inhibitor, and then replacement of growth medium with or without cCr as described in the methods. Quantification data over time is shown for vehicle-treated versus cCr-treated BT549 cells (H, left panel) or for MDA-MB-231 empty vector (EV) or MDA-MB-231 +mCKB cells (H, right panel). Two coverslips were measured simultaneously at each time point across each coverslip/well (n=16 independent images/coverslip/timepoint). Data are representative of three independent experiments. I. Example immunostaining images of invadopodia at experimental endpoint co-stained with cortactin (red) and DAPI (630x magnification, scale bar represents 20 μM). All images were captured with identical laser intensity and exposure settings.

Next, we determined if treatment with cCr inhibits lung colonization, or the growth of pre-established lung micro-metastases. PyMT EV cells were injected into the tail vein and allowed to seed the lungs for 24h. Mice then received daily treatment with cCr at a dose of 1 g/kg or vehicle (0.9% saline, IP) for 21 days. Whereas the surface of lungs from vehicle-treated mice was covered with metastases, very few metastases were visible in the cCr cohort (**Figure 6B**). The majority of mice treated with vehicle developed metastases (7/8, or 87.5%) and 5/8 mice (62.5%) treated with saline developed >100 lesions. In contrast, zero metastases were observed in 5/7 mice in the cCr cohort (71.4%) and 2 mice in each cohort developed fewer than 50 metastases (@^2^, p< 0.05). These results were replicated in an independent experiment, and a third cohort was added in which administration of cCr therapy was delayed by 7 days, a time when lung micro-metastases have formed, such that therapy was then given for a total of 14 days. For mice treated with cCr beginning at day 1, the mean number of metastases was reduced by >10-fold. In contrast, approximately 50% fewer metastases were detected if cCr therapy began on day 7 (**Figure 6C**). There were no detectable macro-metastases in the day 1 cCr cohort, versus 3.8 in the saline group or 1.4 in the day 7 cCr cohort (**Figure 6D**). Over the course of treatment, mice did not develop gross symptoms of toxicity and there was no significant difference in body weight whether animals were treated for either 14 (day 7 cCr) or 21 consecutive days (day 1 cCr) (**Figure 6E**).

### Addition of cyclocreatine to paclitaxel or doxorubicin enhances growth inhibition

In a broad panel of cancer cell lines, cCr repressed cell growth with similar efficacy as conventional agents, including cyclophosamide, doxorubicin, or 5-FU [25,37,38,40]. Rat mammary adenocarcinoma tumor growth was enhanced when cCr (IV, IP) was paired with either cyclophosphamide, doxorubicin, or 5-FU [25], or if the rats were fed chow supplemented with 1% cCr [41]. In pancreatic cancer and in myeloid leukemia, cCr monotherapy inhibits metastasis to distant organs [28,42]. Finally, in HER2+ trastuzumab-sensitive or -resistant breast cancer models, pairing cCr with traszutumab represses cell growth via inhibition of CKMT1 [43]. Based on these observations, we sought to specifically determine in ER-negative breast cancer cell line models if cCr therapy was additive to, or synergistic with, paclitaxel (Taxol) or doxorubicin (DOX), two widely used agents to treat stage IV breast cancer, using a formal isobole testing method [44].

First, we compared CKB expression levels in a panel of breast cancer cell lines representing various molecular subtypes (**Figure 7A** and **Supplemental Figure S4**). MDA-MB-231 and SUM-159 (TNBC) and T47D (ER+, luminal B) cells did not express detectable CKB levels, whereas expression was variable in the other cell lines. CKB protein was most abundant in MDA-MB-453 cells (TNBC), which also express low levels of HER2+ protein, although not above the clinical cutoff for designating HER2-amplification [45]. CKB was moderately expressed in MDA-MB-468 and BT549 TNBC cells (**Figure 7A**), which expressed similar CKB levels as murine PyMT cells (**Figure 1B** and **Figure 7A**). AU565 (HER2+), CAMA1 (ER+, luminal B), HCC2157 (TNBC), and HCC70 (TNBC) also expressed moderate levels of CKB. MDA-MB-436 (TNBC) cells expressed relatively low levels of CKB protein (**Supplemental Figure S4**).

Three TNBC cell line models were selected for evaluation of efficacy of cCr monotherapy versus combination with paclitaxel or DOX, including, MDA-MB-468 (CKB moderate) BT549 (CKB moderate), and MDA-MB-453. (CKB high). BT549 cells also form invadopodia in vitro [46]. The estimated IC_50_ doses of cCr and each chemotherapy were determined by phase confluence assays as described in the methods (**Table 2**). Isobole assays evaluated whether combination therapy was likely to be additive or synergistic (**Table 3**).

**Table 2:**
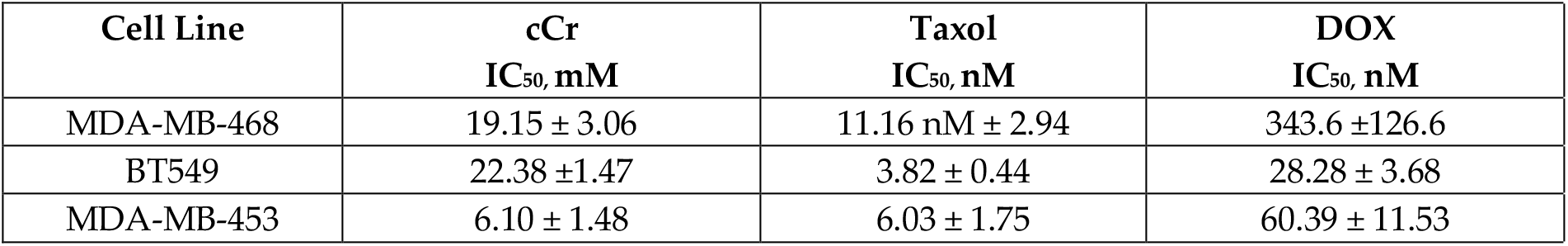
Calculation of monotherapy IC_50_ values in human TNBC cell lines. IC_50_ values ± SEM were determined at 96h as described in the methods. IC_50_ values for independent experiments were calculated using non-linear regression and variable slope fit in Prism 9.0. Results from independent experiments were averaged to report the mean (n=4 replicates/cell line /drug).

| Cell Line | cCr<br>IC<sub>50</sub>, mM | Taxol<br>IC<sub>50</sub>, nM | DOX<br>IC<sub>50</sub>, nM |
| --- | --- | --- | --- |
| MDA-MB-468 | 19.15 ± 3.06 | 11.16 nM ± 2.94 | 343.6 ± 126.6 |
| BT549 | 22.38 ± 1.47 | 3.82 ± 0.44 | 28.28 ± 3.68 |
| MDA-MB-453 | 6.10 ± 1.48 | 6.03 ± 1.75 | 60.39 ± 11.53 |

**Table 3:** Combination indices calculated for human TNBC cell lines pairing cCr with either Taxol or DOX. Isobole studies were performed and data representative of 2 independent isobole assays are shown.

|  | Taxol @ IC<sub>50</sub><br>+ cCr @ IC<sub>30</sub><br>Combination Indices (CI) |  | DOX @ IC<sub>50</sub><br>+ cCr @ IC<sub>30</sub><br>Combination Indices (CI) |  |
| --- | --- | --- | --- | --- |
| Cell Line | IC<sub>50</sub>, TAXOL<br>CI | IC<sub>70</sub>, TAXOL<br>CI | IC<sub>50</sub>, DOX<br>CI | IC<sub>70</sub>, DOX<br>CI |
| MDA-MB-468 | 1.06 | 0.52 | 0.61 | 0.39 |
| BT549 | 0.86 | 0.58 | 1.50 | 1.28 |
| MDA-MB-453 | 0.95 | 2.18 | 0.77 | 1.40 |

In MDA-MB-468 cells, a combination of 10 mM cCr with 25 nM Taxol was synergistic (combination index, CI=0.52), efficiently inhibiting cell growth (**Figure 7B**). For DOX, 100 nM repressed growth by ~50%, but the addition of cCr to DOX further inhibited cell growth (**Figure 7C**); this combination was additive. In contrast, in BT549 cells, cCr was synergistic with a 5 nM dose of Taxol (**Figure 7D**), but no significant change in cell growth was observed when cCr was paired with DOX; in fact, isobole studies suggested antagonistic effects of this combination (CI>1.0) (**Figure 7E, Table 3**). In MDA-MB-453 cells, the most sensitive model to cCr (**Table 2**), the combination of either Taxol (**Figure 7F**) or DOX (**Figure 7G**) with cCr was synergistic at the IC_50_ doses of either agent (CI=0.95 and 0.77, respectively). It has been previously reported that combining cCr with Taxol enhances microtubule stability, leading to the synergistic killing of MCF-7 (ER-positive) cells [47]. To establish if this effect is also observed in ER-negative cells, MDA-MB-468 cells were immunostained for alpha-tubulin. Treatment with cCr slightly disrupted the microtubule network and Taxol stabilized microtubules, but the addition of cCr to Taxol further stabilized tubulin networks, indicating synergy between cCr and Taxol (**Supplementary Figure S5**). The molecular mechanisms of the observed additive effects of cCr in combination with DOX require further investigation.

### Invadopodia formation in TNBC cells depends on creatine kinase activity

To determine if CKB regulates local invasion through the formation of invadopodia, two human TNBC models were used, MDA-MB-231 and BT549 cells. Invadopodia are characterized by focal degradation of a fluorescent gelatin coating deposited onto glass coverslips prior to cell seeding, in addition to the presence of cortactin [48]. We optimized protocols to image loss of gelatin by imaging invadopodia assay coverslips submerged into 24-well dishes using a live-cell imager (**Figure 7H**). Representative experiments of two independent, simultaneously imaged coverslips revealed that treatment of BT549 cells with cCr inhibited loss of gelatin over time (52h), whereas ectopic expression of CKB in MDA-MB-231 cells stimulated gelatin loss over 24h (**Figure 7H**). The presence of cortactin in areas of invadopodia formation was validated by immunofluorescence at the end of the imaging (**Figure 7I**; BT549 images in **Supplementary Figure S6A**.). We also immunostained for CKB in endpoint invadopodia in MDA-MB-231 +mCKB cells. Representative images comparing empty vector and +mCKB cells revealed CKB protein is detectable at the edges of invadopodia (**Supplementary Figure S6B**).

## Discussion

Although multiple roles for HIF-1α in mediating breast cancer phenotypes are well-defined [49], many individual HIF-1α-dependent genes remain to be characterized for their role in breast tumor progression and metastasis in specific breast cancer subtypes. We identified CKB as a HIF-1α-dependent target gene necessary for tumor outgrowth and lung metastasis in the MMTV-PyMT transgenic model of MBC. Higher *CKB* mRNA or protein levels were also found to be prognostic of reduced relapse-free survival (RFS) across all molecular subtypes of breast cancer. The likelihood of RFS was further decreased for patients diagnosed with basal-like breast cancers. Since basal-like and HER2-enriched subtypes exhibit high constitutive expression of HIF-1α and HIF-1 target genes [50,51], we focused on understanding the function of CKB in ER-negative models.

HIF-dependent, but not hypoxia stimulus-dependent, regulation of *Ckb* mRNA and protein levels was observed in PyMT cells. Since HIF protein stabilization can be induced independently of hypoxia by several growth factor signaling pathways, including EGF and HER2 [52], the HIF-1-dependent, but not hypoxia-induced, expression of CKB in TNBC may be mediated in this manner. Cell stressors such as low glucose, change in pH, and the production of reactive oxygen species (ROS) all increase HIF-1α expression [53], and could up-regulate CKB expression independent of hypoxia. In contrast, in ER+ MCF-7 breast cancer cells, *CKB* mRNA levels increased in response to hypoxia. In both models, ChIP assays demonstrated that CKB is a direct HIF transcriptional target in the breast epithelium.

CKB mRNA and protein levels were also decreased in HIF-1 KO end-stage, whole tumors, although the reduction was modest relative to cultured cells. A significant difference in *Ckb* mRNA levels in tumors was observed only after normalization to the epithelial marker K18. Since HIF-1 loss, and therefore, reduced CKB, is targeted only to the tumor epithelium, a significant proportion of the CKB signal observed in whole tumor extracts is likely derived from the stroma. CKB is expressed in adipose, macrophages, and endothelial cells [29,54–56].

The biological activities predominantly impacted in response to *Ckb* shRNA KD were related to metastatic potential, including repression of invasion through extracellular matrix (ECM) and reduced lung metastasis. ATP production was also inhibited. Surprisingly, Seahorse analysis revealed decreases in both the glycolytic and mitochondrial respiration (OXPHOS) arms of cellular metabolism, with the most prominent changes observed for sh61 shRNA cells. Glycolytic capacity was reduced by 53%, basal respiration was reduced by 46% and maximal respiration was reduced by 55%. Further, the basal and maximum respiration rates of sh61 KD cells were similar (59.9 vs. 63.6 mpH/min/RFU), suggesting an inability to respond to increased demand for energy. We conclude that whereas either cellular proliferation or cell motility can tolerate decreased ATP production when CKB levels are low, the creatine kinase phosphagen arm is essential to produce the energy necessary for cell invasion and metastatic colonization. In other breast cancer cell types that regulate CKB in a hypoxia-dependent manner, like MCF-7 cells, increased CKB expression may be necessary to supplement energy production when oxygen is limiting and glycolysis is the predominant pathway to generate ATP. It has been suggested that cytosolic CKs interact with the mitochondrial CK isoforms to maintain energy flux [15]. In thermogenic brown fat, CKB directly shuttles between mitochondria and the cytoplasm, directly regulating mitochondrial ATP turnover [56]. CKB deletion in the presence of oxygen impaired energy production through glycolysis and mitochondrial respiration, similar to our observations in ER-negative breast cancers.

Despite minimal changes in proliferation *in vitro* in response to loss of CKB, there was a robust decrease in PyMT *Ckb* KD tumor growth rate. Remarkably, deletion of this single HIF-1 target gene produced tumors with similar volumes to HIF-1 KO cells, suggesting that tumor cell-intrinsic CKB directly influences the local surrounding microenvironment to promote tumorigenesis. It is possible that loss of epithelial-derived CKB disrupts a paracrine network necessary to promote tumor growth. For example, local extracellular secretion of CKB by colon cancer cells into the liver microenvironment promotes metastatic outgrowth through extracellular production of phosphocreatine (PCr), which was then imported into tumor cells as an energy source [27]. In this model, implantation of an osmotic pump releasing cCr into peritoneum also repressed liver metastasis [27].

Tail vein assays revealed a significant decrease in lung colonization either in response to *Ckb* KD or by inhibiting CK activity with daily cCr systemic treatment. cCr significantly inhibited formation of lung macro-metastases and the conversion of pre-established micro-metastases to macro-metastases. Because overall survival increases when *Ckb* is deleted in tumors re-generated from MTECs orthotopically implanted into the WT mammary fat pad, it is also clear that CKB is necessary for efficient completion of the entire metastatic cascade, beginning with local invasion. This conclusion is also supported by our observations that loss of CKB in PyMT cells impairs invasion whereas over-expression of CKB in MDA-MB-231 cells enhances not only chemotaxis and cell invasion, but also promotes invadopodia activity. The apparent relationship between HIF-1, a master regulator of cellular metabolism, and CKB, and the degradation of extracellular matrix by invadopodia is intriguing. In a proteomics screen to identify factors enriched in invadopodia, multiple HIF-dependent metabolic enzymes were identified, including GAPDH, enolase 1, lactate dehydrogenase (LDH), pyruvate kinase, muscle 2 (PKM2), and phosphoglycerate kinase 1 (PGK1) [57]. Additional studies are warranted to ascertain how HIF-1α-dependent modulation of CKB expression contributes to each step of the metastatic cascade. Of note, creatine supplementation, which increases creatine kinase flux, is not sufficient to change primary tumor growth rate, but significantly enhances lung metastasis [58]. Overall, tight regulation of cellular metabolism by CKs is likely critical for a cell’s ability to invade.

Local ATP generation coordinated by CKB to facility cell motility was previously suggested for astrocytes and fibroblasts [29]. The direct relationship of creatine flux to ATP levels and to cell motility was recently revealed using pancreatic cancer models [28]. In this study, CKB, but not other CK isoforms, was shown to be a Yes-associated protein (YAP)-responsive mechanosensory responder. In particular, YAP increased CKB expression and CK activity in response to a stiff extracellular environment. Increased expression of YAP promotes breast tumorigenesis, although it is dispensable for normal mammary gland development [59]. Connections between YAP and HIF and CKB are of interest since YAP can induce HIF-1α [59] through mechanical loading [60], and since hypoxia/HIF-1 can also stimulate YAP activity through 14-3-3-zeta [61], potentially setting up feed-forward loops executed via CKB that are necessary for metastatic potential.

Cytotoxic chemotherapies are frontline treatments for stage IV TNBC. We observed that pairing cCr with either paclitaxel or doxorubicin enhanced cCr efficacy. The most potent inhibition of cell growth occurred when cCr was paired with paclitaxel. cCr treatment was also synergistic with DOX near the IC_50_ of DOX in two TNBC cell lines (MDA-MB-468 and MDA-MB-453), but was antagonistic to cCr in BT549 cells. Interestingly, doxorubicin can impact heart microtubule structure by impairing reassembly [62]. Down-regulation of CKB in Skov3 ovarian cancer cells was shown to enhance sensitivity to doxorubicin [20]. However, the exact mechanisms of DOX synergy with cCr remain undefined. We conclude that cCr therapy inhibits growth and impairs chemotaxis/cell motility through both its effects on cellular metabolism and microtubule dynamics.

There is increasing evidence from a variety of solid cancers that CK inhibition is a promising clinical intervention for patients with metastatic disease. In addition, cCr likely crosses the blood-brain barrier and phosphorylated-cCr functions as a phosphagen in the brain [63,64], suggesting that creatine analogs may effectively target brain metastases. Systemic cCr therapy is well-tolerated in rodents [65] and in limited clinical trials [15]. In addition, deletion of *Ckb* or systemic administration of cCr is protective against bone loss in mice [66], suggesting additional potential clinical applications to prevent osteolytic destruction of bone, the most common site of metastasis in breast cancer patients. A new generation derivative of cCr known as LUM-001 with enhanced bio-availability was developed to treat creatine transporter deficiency. LUM-001 was tested in rodents to treat neurodevelopmental cognitive disorders, including autism [63]. Clinical trials using LUM-001 are in enrollment (www.clinicaltrials.gov; NCT02931682) after pre-clinical studies were completed by the NIH Therapeutics for Rare and Neglected Diseases (TRND) program [67].

## Conclusions

In summary, our results demonstrate that CKB is a major effector of HIF-1-mediated promotion of metastatic phenotypes in ER-negative breast cancer *in vitro* and *in vivo*. CKB is capable of driving aggressive phenotypes in the context of normoxia or hypoxia, likely through its combined roles in managing cellular metabolism, cell cycle progression and microtubule dynamics. A growing body of evidence indicates that cCr has anti-tumor efficacy in multiple breast cancer subtypes, including HER2+ breast cancers, and as reported herein, in ER-negative/basal breast cancer models. New clinical trials to explore cCr monotherapy versus combination with either Taxol or doxorubicin would be predicted to demonstrate prolonged survival through reduced survival of cancer cells into the circulation, during colonization, and by impairing the growth of cells already disseminated to distant organs, including lung, brain and bone metastases.

## Supporting information

SuppFigures and Legends

## Author Contributions

R.I. Krutilina: Methodology development, investigation, writing, including review and editing. H.C. Playa: Methodology development, investigation, writing, including review and editing. The first two authors contributed equally and are listed in alphabetical order. D.L. Brooks: Conceptualization, methodology development, investigation, writing, including review and editing. L.P. Schwab: Methodology development, investigation, and writing-editing. D.N. Parke: Methodology development, investigation. D. Oluwalana: Investigation. D. Layman: Investigation. M. Fan and DL. Johnson: Investigation and resources related to data analysis and bioinformatics support. DJ. Yue: Generation of research materials and methodology. H. Smallwood: Methodology and investigation. T.N. Seagroves: Conceptualization, supervision, funding acquisition, investigation, writing-original draft, and writing-review and editing.

## Institutional Review Board and IACUC Statement

All studies were reviewed to be Not Human Subjects Research by the UTHSC Institutional Review Board (IRB); study ID, 0705002NHSR. All animal studies were consistent with U.S. Public Health Service policies and the Guide for the Care and Use of Laboratory Animals. Animal protocols were approved by the local UTHSC Institutional Animal Care and Use Committee (Study investigator: T. Seagroves).

## Data Availability Statement

Microarray data are publicly available through the Gene Expression Omnibus (GSE#183694, released September 10, 2021).

## Acknowledgments

We are grateful to the staff of the institutional core facilities at The University of Tennessee Health Science Center who supported this work, including Dr. William Taylor and Mr. Lorne Rose of the Molecular Resource Center (MRC). Dr. Deidre Daria assisted with cell cycle analysis at the FCCS Core, and Dr. Daniel Johnson provided optimized data analysis workflows and long-term data storage at the Molecular Bioinformatics (mBIO) core. All paraffin-embedded tissue sections were prepared by the Research Histology Core (RHC) with support from LaShawn Barnett. We thank the Department of Pathology and the Cancer Center at UTHSC for support to digitally scan all immunostained slides and for providing access to the 3D Histech analysis workstation and the Cancer Center for support of the IncuCyte S3 live-imaging system. The UTHSC Department of Pediatrics supports the Seahorse XF instrument managed by Dr. Heather Smallwood. We thank Dr. Athena Starland-Davenport and Dr. Rob Williams for providing access to the Nikon ECLIPSE Ti2 microscope.

## Supplementary Materials and Methods

### Chemicals

Cyclocreatine (cCr) and doxorubicin was purchased from either Sigma-Aldrich (#377627, or #D1515; 98% purity) or Cayman Chemical Company (cat# 20649, Ann Arbor, MI, >95% purity), and paclitaxel (>99.5% purity) was purchased from LC Laboratories (cat.# P-9600, Woburn, MA). cCr stock was freshly diluted to 50 mM in complete growth medium immediately before use. Paclitaxel (Taxol) and doxorubicin (DOX) were prepared in DMSO as 20 mM and 5 μM stocks, respectively, and stored at −20°C; thawed stocks were serially diluted in DMSO to prepare working stocks for *in vitro* assays. For use *in vivo*, cCr was dissolved to ~1.92 mg/mL into sterile 0.9% saline using gentle heat, sterile filtered, and at −20°C. Prior to treatment, the frozen aliquot was heated to ~50°C, transferred to a sterile glass evacuated vial, and stored protected from light in a thermos heated to 37°C. cCr was immediately injected into mice (IP) at a final dose of 1.0 g/kg/day.

### RNA preparation and harvest for microarray profiling

HIF-1 WT and KO PyMT cells grown in DMEM/F12 medium (n=6 wells/genotype) supplemented with 2% FBS and 15 mM HEPES. At ~80% confluence, plates were left at normoxia or transferred to hypoxia (0.5% O_2_) for 6h, when HIF-1 transcriptional activity peaks [1]. At harvest, individual wells were washed with cold PBS, scraped, and cells pelleted and flash frozen. Total RNA was prepared from partially thawed cell pellets using RNABee reagent (Tel-Test, Friendswood, TX). Independent replicate total RNA samples (n=3) were hybridized to the MouseRef-6v1.1 Expression BeadChip Kit (Illumina, San Diego, CA) at the UTHSC Molecular Resource Center of Excellence (MRC). Supplementary Tables S1-S3 include differentially-expressed gene lists from the following comparisons, respectively: Table S1: HIF-1 WT @normoxia vs. HIF-1 WT @hypoxia, Table S2: HIF-1 WT vs. HIF-1 KO, @normoxia and Table S3: HIF-1 WT vs. HIF-1 KO, @hypoxia.

### Real-time quantitative PCR (qPCR)

Total RNA isolated using the RNABee reagent was converted to cDNA using the High-Capacity cDNA Reverse Transcription Kit (Applied Biosystems, Waltham, MA). qPCR was performed on the Roche LC480 instrument using default cycling parameters. Crossing point (Cp) values were normalized based on the expression of the integrator complex subunit 3 (*Ints3*) for mouse genes or cyclophilin A (*PPIA*) for human genes. To compensate for any changes in epithelial content in total RNA extracted from whole tumors, since only the tumor epithelium is deleted for *Hif1a*, *Ints3*-normalized Cp values were also normalized to *Krt18* (K18).

### Immunofluorescence staining

Cells were cultured to sub-confluence in multi-well chamber slides (ibidi, Gräfelfing, Germany). At harvest, cells were washed with cold PBS and fixed with 4% paraformaldehyde/PBS before permeabilization with 0.1% Triton-X, washing in PBS + Tween-20 (PBST), and blocking with buffer containing both BSA and normal serum. Primary antibodies to CKB, cortactin, or alpha-tubulin were incubated at 4°C overnight in blocking buffer. Cortactin and tubulin were detected with anti-IgG secondary antibodies for 1h at RT (either AlexaFlour-488, or −594, Life Technologies). Tertiary amplification was used to detect CKB; after primary antibody, cells were incubated with donkey anti-rabbit-biotin-X-IgG secondary antibody (cat. #A16027, Life Technologies, 1;400), washed, and then incubated with Alexa-Fluor-594-conjugated streptavidin. Prior to mounting, cells were counterstained with DAPI, washed and then ibidi mounting medium was added.

### Protein extraction and western blotting

Whole cell extracts (WCE) and high-salt enriched WCE (HS-WCE) fractions were prepared as in [1]. To detect HIF-1α, HS-WCE (5 μg) was resolved on 3-8% Tris-Acetate gels. For CKB, WCE (20-40 μg input) were resolved on either 4-12% Tris-Bis or 10% Tris-Bis BOLT gels (Life Technologies) and transferred onto PVDF Fluorescent (PVDF-FL) membrane (Millipore, Burlington, MA). To confirm equivalent loading, membranes were immunostained with antibodies as described in the figure legends or were stained prior to blotting with Ponceau S (Sigma, St. Louis, MO) and scanned. Ponceau S-stained eembranes were de-stained prior to blocking with either 5% non-fat dry milk (NFDM)/TBST for enhanced chemiluminescence (ECL) or with Odyssey blocking buffer (LiCor Biosciences) for near-infrared (NIR) detection. Membranes were probed with primary antibodies followed by extensive washing and incubation with anti-rabbit whole IgG secondary antibodies conjugated to either HRP (Jackson Immunologicals, West Grove, PA) or to NIR fluorophores (LiCor Biosciences, Lincoln, NE). Antibody complexes were detected by ECL and membranes exposed to film, or membranes were directly imaged using the LiCor Odyssey or Azure Sapphire NIR imaging systems. In some cases, membranes were stripped with ReBlot Plus Mild (Millipore), then re-blocked and incubated with antibodies to ß-tubulin or PARP. Alternatively, .tiff images of Ponceau-S-stained membranes were analyzed by ImageStudio densitometry analysis (LiCor Biosystems) to agnostically quantitate whole lane signal.

### Patient database mining

Datasets were analyzed using The Cancer Genome Atlas (TCGA) database [2] and Kaplan-Meier (KM)Plotter [3]. The level of *CKB* mRNA expression in breast tumor subtypes was derived from the TCGA data portal (http://cancergenome.nih.gov/). Level three normalized data derived from Illumina RNASeqv2 data compared mRNA expression, and samples were stratified by tumor subtype based on the PAM50 method (last accessed October 4, 2016). Plots with *p*-values ≤ 0.05 (ANOVA with Bonferroni correction) were considered significant. For regression-free survival (RFS) analysis, KMPlotter was stratified by CKB (200884_at) mRNA or protein (P12277) expression in breast cancer patients (last accessed February 3, 2021).

### Promoter analysis and Chromatin immunoprecipitation (ChIP) assays

The mouse and human *CKB* proximal promoters (−2000 to +500 bp) were scanned for putative functional hypoxic response elements (HREs) using the Transcription Factor Matrix (TFM) Explorer algorithm and weight matrices available from JASPER and TRANSFAC. PyMT HIF-1 WT and KO cells and MCF7 EV (pLKO.1-puro) or shHIF1A transduced cells described in [4] were cultured at normoxia or 0.5% O_2_ (hypoxia) for 6-24h and fixed with 1% formaldehyde for 12min. Positive controls included a previously validated functional HRE in the *Vegf* promoter for PyMT cells [5] and a previously validated functional HRE identified in the *EPO* promoter [6] for MCF-7 cells.

### PyMT Ckb shRNA knockdown (KD) cell lines

PyMT HIF-1 WT cells were used to create *Ckb* stable shRNA knockdown (KD) via lentiviral transduction. First, a *Ckb* shRNA library containing 3 independent *Ckb* targeting sequences individually cloned into the pLKO.1-puro vector was purchased from Open Biosystems (clone IDs: TRCN00000246<u>59</u>, “sh59”, TRCN00000264<u>60</u>, “sh60” and TRCN00000246<u>61</u>, “sh61”; Supplementary Table S7). Cells (1×10^6^) were transfected with each shRNA plasmid (4 μg) by nucleofection with kit T and program setting 24 (Nucleofector, Lonza). Lentivirus particles for pLKO.1-empty (empty vector, EV), and the pLKO.1-puro sh59 and sh61 constructs were produced by Dr. Junming Yue of the UTHSC Viral Vector Core. PyMT cells were transduced with lentivirus particles (MOI=50). Puromycin (2 μg/mL) was added 72h later and a “pool” of surviving clones (representing several hundred colonies) was established for each shRNA. Stably transduced lines were maintained in growth media plus 1 μg/mL puromycin; antibiotic selection was removed from cells at least 4 days before biological assays or treatments with cCr.

### Human breast cancer cell lines, mycoplasma testing and cell line authentication

All cells were obtained from the Fan laboratory at UTHSC via the American Type Culture Collection (ATCC) and grown in base media (DMEM-Hi: MDA-MB-231 and MDA-MB-453 cells, DMEM/F-12: MDA-MB-468 cells or RPMI: BT549 cells) supplemented with 10% FBS (cat.# FB-01, Omega Scientific, Tarzana, CA). All cells were authenticated at the University of Arizona Genetics Core. Cells were routinely screened for mycoplasma using the Lonza MycoAlert kit.

### PyMT cell proliferation assay by WST-1

Cells were grown at normoxia or hypoxia in medium buffered with 15 mM HEPES. The day before enumeration, 20,000 cells were plated into 96-well plates in normal growth medium (n=3 wells/genotype/oxygen tension/time point). The medium was changed 24h post-plating, when the first cell harvest was collected (t=0h time point), but medium was not replenished for the duration of the experiment. Background absorbance was subtracted and raw data normalized to the EV controls (% maximal proliferation vs. EV). To generate a cCr dose-response curve, at 24h post-seeding, growth media without drug or with media containing increasing doses of cCr (from 2 mM to 50 mM) was replaced. Cells were incubated for 96h and then analyzed by the WST-1 assay (Chemicon, cat.#2210, Burlington, MA). Growth inhibition was measured as the change in absorbance over time, normalized to the vehicle control for each cell line (set to 100%).

### Wound healing assays

PyMT EV, HIF-1 KO, and *Ckb* KD cells were plated in 12-well plates in growth medium (30,000 cells/well; n=3 wells per genotype) and grown to 100% confluence. A scratch with a 1 mL tip in a vertical and horizontal direction was applied to each well. Cells were washed to remove the detached cells and fresh growth medium added (t=0h). Images were captured at 100x magnification at 0, 8, 24, 32, 48, 56, and 72h and analyzed for open wound area/total area using ImageJ. Additional wound healing assays were performed using the IncuCyte S3 live-cell imaging system. PyMT EV cells were pre-incubated for 24h with 25 mM cCr in growth media supplemented with 2% FBS. Cells were plated into 96-well format Image Lock microplates (Sartorius, Göttingen, Germany) at 40,000/cells/well (n=8 technical replicates/condition) such that cells would be 100% confluent approximately 24h later. The WoundMaker tool created a uniform 700-μM scratch/well, the wounds were washed and growth medium containing either vehicle or 25 mM cCr was applied. Wounded EV cells that were not pre-treated with cCr were exposed to vehicle or to cCr (25 mM). Raw data were analyzed using the total wound area algorithm and exported into Prism 9.0.

### Invasion assays

PyMT EV, HIF-1 KO, and *Ckb* KD cells were first gradually weaned from growth medium supplemented with 2% FBS to 0.5% FBS as in [1]. Cells were then cultured 18h in serum-free DMEM/F-12 medium. The next day, 25,000 cells were plated onto control inserts or Matrigel-coated transwell inserts (BD Biosciences, San Jose, CA) and attracted to wells containing DMEM/F12 medium supplemented with 5% FBS (n=3 wells/genotype). The mean cell invasion index corrected for random migration was calculated at 48h post-seeding. To determine if re-expression of CKB in HIF-1 KO PyMT cells would rescue invasion, vector control, or HIF-1 KO cells stably transfected to express mCKB were gradually weaned to 0.5% FBS and then plated at a density of 40,000 cells into transwell inserts (with and without coating with Matrigel) and attracted to 5% FBS for 48h. MDA-MB-231 transiently transfected cells (empty vector or +mCKB) were serum-starved overnight before plating onto transwell inserts and then attracted to medium supplemented with 10% FBS for 24h. For studies comparing normoxia to hypoxia, changes in PyMT cell invasion are expressed as the fold change relative to the invasion index observed for HIF-1 WT cells cultured at normoxia (fold change = 1.0). Otherwise, all other data are expressed as the % of invading cells (invading cells/total cells plated).

### Intracellular ATP Assay

Intracellular ATP levels were compared using the high sensitivity ATP Bioluminescence Assay HS II kit (cat. #11699709001, Sigma). PyMT cells were grown to 80% confluence prior to incubation at normoxia or hypoxia for 24h (n=3 per genotype/condition). Cells were harvested and lysed using the kit lysis reagent supplemented with cOmplete EDTA-free tablets (Sigma, cat#4693132001). Luciferase reagent was added 1s prior to a 10s integrated reading on a single-tube luminometer. A blank (reagent only) reading was obtained and subtracted from all sample readings. An ATP standard curve was generated to plot bioluminescence versus molarity on a log-log scale and used to calculate intracellular ATP concentrations. DNA concentrations for each individual sample were measured using a Qubit® fluorometer (Life Technologies), and ATP concentrations was normalized to total DNA.

### Chemotaxis assays

MDA-MB-231-NR cells transiently transfected (48h) with empty vector (EV, pCMV-6-Entry) or expressing mCKB (+mCKB) were seeded at a density of 1,000 cells/per well of a 96-well ClearView chemotaxis plate (cat. #4582, Sartorius) (n=4-6 technical replicates/cell line), which allows cell tracking in real-time through optically clear membrane inserts. Cells were exposed to a reservoir containing DMEM +10% FBS as the chemoattractant. The chemotaxis analysis software tool was used to quantify cell migration. Raw data were normalized for initial plating density prior to export to Prism 9.0.

### Cytotoxicity assays

PyMT cells were seeded at 5,000 cells per well in 96-well dishes and allowed to adhere overnight. The following day, growth medium (vehicle control) or cCr was added. A 10 μM working stock solution of CytoTox Green reagent (Sartorius, cat.# 4633) was made fresh and added to each well at a final concentration of 250 nM. The plate was placed into the IncuCyte S3 instrument and cells were allowed to equilibrate for approximately 1h before initiation of imaging; each well was imaged in three locations every 4h for a total of 48h. At the experimental endpoint, the phase and fluorescent masking software algorithms were applied. The data were exported to Prism 9.0 and normalized to corresponding initial seeding density measured by phase contrast. Data are representative of two independent biological experiments with n=6 technical replicates/time point/cell line.

### Cell cycle analysis of PyMT cells

PyMT EV or sh61 KD cells were plated into 6-well dishes and when sub-confluent were treated with vehicle control or with 25 mM cCr for either 24h (added 24h prior to experiment endpoint) or 48h. Harvested cell pellets were washed twice with cold PBS and re-suspended in 5mL ice-cold 70% ethanol prior to storage at 4°C. Prior to cytometry, samples were washed with PBS and treated with RNaseI. Setting aside an unstained sample as the gating control, propidium iodide [50 μg/mL] was added and incubated at 37°C for 15min prior to analysis on a BioRad ZE5 flow cytometer at the UTHSC Flow Cytometry and Cell Sorting (FCCS) Core. Raw data were exported to ModFit for analysis (n=3 replicates/genotype/time point, and at least duplicate biological replicates per time point).

### Tissue immunohistochemistry and immunostaining quantification

Harvested tumors were bisected with a razor and fixed for 8-24h in 10% neutral-buffered formalin (NBF), followed by paraffin embedding and sectioning. Antigen retrieval was performed in 1x citrate buffer (pH 6.0) in a pressure cooker. Endogenous peroxidase was blocked by 3% H_2_0_2_/MeOH for 20 minutes, followed by washing in water then PBS. All slides were blocked in 10% normal serum/PBST at least 1h at RT before incubation of primary antibody overnight in humid chambers at 4°C. Slides were washed then stained with the ABC Elite anti-rabbit kit (cat. # PK-6100, Vector Labs, Burlingame, CA) and developed using DAB Impact (cat. #SK-4105, Vector Labs) and counterstained in hematoxylin (cat.# H3401, Vector Labs). Whole slides were digitally scanned using a 3DHISTECH PANORAMIC slide digitizer (3D Histech, Ltd. software, Budapest, Hungary). Staining intensity was analyzed by pixel counts using built-in densitometry algorithms for nuclear or cytoplasmic localization. Following masking to exclude necrosis, whole tumors were analyzed after the general background was set using an adjacent tissue section developed without primary antibody. Antibody reagents are listed in Supplementary Table S5.

### Invadopodia assays using human TNBC cells

Glass coverslips were coated with OregonGreen-gelatin (cat. #G13186, ThermoFisher) as in [7]. Cells (BT549, 12,000/well or MDA-MB-231, 15,000/well) were seeded onto coverslips in growth media containing a reversible MMP inhibitor, batimistat (10μM; cat. # SML0041, Sigma) and incubated overnight. The next day, the media was replaced with fresh growth media (vehicle) or cCr (25 mM) was also added. 12-well plates containing seeded coverslips were placed into the IncuCyte S3 imager and simultaneously imaged for green fluorescence and phase contrast (n=16 images/coverslip/time point). Green fluorescent area was normalized to seeding density per image and graphed as a ratio of change over time in Prism 9.0. At the end of the imaging, optimized per cell line, coverslips were fixed and immunostained prior to mounting directly onto glass slides using ProLong Gold Antifade media (cat.# P36930, ThermoFisher). Coverslips were imaged on a Nikon ECLIPSE Ti2 microscope as described.

### Growth inhibition on TNBC cell lines during chemotherapy treatment

One day before measuring growth, cells were seeded into 96-well flat-bottomed plates such that assay imaging would begin the next day at ~20% confluence (MDA-MB-468: 8,000 cells/well; MDA-MB-453: 12,000 cells/well; MDA-MB-231 or BT549 cells: 5,000 cells/well). At t= − 1h, growth media from seeded cells was removed and each dilution of chemotherapy drug, or the vehicle control, was spiked into fresh media containing either IncuCyte Nuclight Rapid Red reagent (cat.# 4717, Sartorius) or Miami Green (cat.# EMI001, Kerafast, Boston, MA) to enumerate cell nuclei. Plates were imaged in the IncuCyte S3 instrument. Growth inhibition percentage was calculated relative to vehicle controls, as calculated by measurement of red or green fluorescent units localized to the nucleus after applying the IncuCyte masking algorithm to enumerate cell count. Data were exported to Prism 9.0 and normalized to initial seeding density. Drug synergism was calculated using the isobole method [8], wherein a combination index of >1.0 indicates antagonism, an index of ~1.0 is additive and an index of <1.0 suggests synergism.

## References

1. National Cancer Institute Surveillance, E., and End Results (SEER) Program. Cancer Stat Facts: Female Breast Cancer. Available online: https://seer.cancer.gov/statfacts/html/breast.html (accessed on August 16, 2021).

2. Bailey, K.M.; Wojtkowiak, J.W.; Hashim, A.I.; Gillies, R.J. Targeting the metabolic microenvironment of tumors. Adv Pharmacol 2012, 65, 63–107, doi:10.1016/B978-0-12-397927-8.00004-X.

3. Zhao, Y.; Butler, E.B.; Tan, M. Targeting cellular metabolism to improve cancer therapeutics. Cell Death Dis 2013, 4, e532, doi:10.1038/cddis.2013.60.

4. Dang, N.H.; Singla, A.K.; Mackay, E.M.; Jirik, F.R.; Weljie, A.M. Targeted Cancer Therapeutics: Biosynthetic and Energetic Pathways Characterized by Metabolomics and the Interplay with Key Cancer Regulatory Factors. Curr Pharm Des 2013.

5. Semenza, G.L. HIF-1 mediates metabolic responses to intratumoral hypoxia and oncogenic mutations. J Clin Invest 2013, 123, 3664–3671, doi:10.1172/JCI67230.

6. Chiche, J.; Brahimi-Horn, M.C.; Pouyssegur, J. Tumour hypoxia induces a metabolic shift causing acidosis: a common feature in cancer. J Cell Mol Med 2010, 14, 771–794, doi:10.1111/j.1582-4934.2009.00994.x.

7. Rohwer, N.; Cramer, T. Hypoxia-mediated drug resistance: novel insights on the functional interaction of HIFs and cell death pathways. Drug Resist Updat 2011, 14, 191–201, doi:10.1016/j.drup.2011.03.001.

8. Brown, J.M. Exploiting the hypoxic cancer cell: mechanisms and therapeutic strategies. Mol Med Today 2000, 6, 157–162.

9. Schwab, L.P.; Peacock, D.L.; Majumdar, D.; Ingels, J.F.; Jensen, L.C.; Smith, K.D.; Cushing, R.C.; Seagroves, T.N. Hypoxia-inducible factor 1alpha promotes primary tumor growth and tumor-initiating cell activity in breast cancer. Breast Cancer Res 2012, 14, R6, doi:10.1186/bcr3087.

10. Xia, Y.; Choi, H.K.; Lee, K. Recent advances in hypoxia-inducible factor (HIF)-1 inhibitors. Eur J Med Chem 2012, 49, 24–40, doi:10.1016/j.ejmech.2012.01.033.

11. Doedens, A.; Johnson, R.S. Transgenic models to understand hypoxia-inducible factor function. Methods Enzymol 2007, 435, 87–105, doi:10.1016/S0076-6879(07)35005-2.

12. Keith, B.; Johnson, R.S.; Simon, M.C. HIF1alpha and HIF2alpha: sibling rivalry in hypoxic tumour growth and progression. Nat Rev Cancer 2012, 12, 9–22, doi:10.1038/nrc3183.

13. Wallimann, T.; Wyss, M.; Brdiczka, D.; Nicolay, K.; Eppenberger, H.M. Intracellular compartmentation, structure and function of creatine kinase isoenzymes in tissues with high and fluctuating energy demands: the ‘phosphocreatine circuit’ for cellular energy homeostasis. Biochem J 1992, 281 ( Pt 1), 21–40.

14. Wallimann, T.; Tokarska-Schlattner, M.; Schlattner, U. The creatine kinase system and pleiotropic effects of creatine. Amino Acids 2011, 40, 1271–1296, doi:10.1007/s00726-011-0877-3.

15. Wyss, M.; Kaddurah-Daouk, R. Creatine and creatinine metabolism. Physiol Rev 2000, 80, 1107–1213.

16. Glover, L.E.; Bowers, B.E.; Saeedi, B.; Ehrentraut, S.F.; Campbell, E.L.; Bayless, A.J.; Dobrinskikh, E.; Kendrick, A.A.; Kelly, C.J.; Burgess, A.; et al. Control of creatine metabolism by HIF is an endogenous mechanism of barrier regulation in colitis. Proc Natl Acad Sci U S A 2013, 110, 19820–19825, doi:10.1073/pnas.1302840110.

17. Streijger, F.; Oerlemans, F.; Ellenbroek, B.A.; Jost, C.R.; Wieringa, B.; Van der Zee, C.E. Structural and behavioural consequences of double deficiency for creatine kinases BCK and UbCKmit. Behav Brain Res 2005, 157, 219–234, doi:10.1016/j.bbr.2004.07.002.

18. Mooney, S.M.; Rajagopalan, K.; Williams, B.H.; Zeng, Y.; Christudass, C.S.; Li, Y.; Yin, B.; Kulkarni, P.; Getzenberg, R.H. Creatine kinase brain overexpression protects colorectal cells from various metabolic and non-metabolic stresses. J Cell Biochem 2011, 112, 1066–1075, doi:10.1002/jcb.23020.

19. Zarghami, N.; Giai, M.; Yu, H.; Roagna, R.; Ponzone, R.; Katsaros, D.; Sismondi, P.; Diamandis, E.P. Creatine kinase BB isoenzyme levels in tumour cytosols and survival of breast cancer patients. Br J Cancer 1996, 73, 386–390.

20. Li, X.H.; Chen, X.J.; Ou, W.B.; Zhang, Q.; Lv, Z.R.; Zhan, Y.; Ma, L.; Huang, T.; Yan, Y.B.; Zhou, H.M. Knockdown of creatine kinase B inhibits ovarian cancer progression by decreasing glycolysis. Int J Biochem Cell Biol 2013, 45, 979–986, doi:10.1016/j.biocel.2013.02.003.

21. Chen, H.; Pimienta, G.; Gu, Y.; Sun, X.; Hu, J.; Kim, M.S.; Chaerkady, R.; Gucek, M.; Cole, R.N.; Sukumar, S.; et al. Proteomic characterization of Her2/neu-overexpressing breast cancer cells. Proteomics 2010, 10, 3800–3810, doi:10.1002/pmic.201000297.

22. Chen, W.Z.; Pang, B.; Yang, B.; Zhou, J.G.; Sun, Y.H. Differential proteome analysis of conditioned medium of BPH-1 and LNCaP cells. Chin Med J (Engl) 2011, 124, 3806–3809.

23. Xu, Y.; Cao, L.Q.; Jin, L.Y.; Chen, Z.C.; Zeng, G.Q.; Tang, C.E.; Li, G.Q.; Duan, C.J.; Peng, F.; Xiao, Z.Q.; et al. Quantitative proteomic study of human lung squamous carcinoma and normal bronchial epithelial acquired by laser capture microdissection. J Biomed Biotechnol 2012, 2012, 510418, doi:10.1155/2012/510418.

24. Zeng, G.Q.; Zhang, P.F.; Deng, X.; Yu, F.L.; Li, C.; Xu, Y.; Yi, H.; Li, M.Y.; Hu, R.; Zuo, J.H.; et al. Identification of candidate biomarkers for early detection of human lung squamous cell cancer by quantitative proteomics. Mol Cell Proteomics 2012, 11, M111 013946, doi:10.1074/mcp.M111.013946.

25. Teicher, B.A.; Menon, K.; Northey, D.; Liu, J.; Kufe, D.W.; Kaddurah-Daouk, R. Cyclocreatine in cancer chemotherapy. Cancer Chemother Pharmacol 1995, 35, 411–416, doi:10.1007/s002800050255.

26. Mulvaney, P.T.; Stracke, M.L.; Nam, S.W.; Woodhouse, E.; O’Keefe, M.; Clair, T.; Liotta, L.A.; Khaddurah-Daouk, R.; Schiffmann, E. Cyclocreatine inhibits stimulated motility in tumor cells possessing creatine kinase. Int J Cancer 1998, 78, 46–52.

27. Loo, J.M.; Scherl, A.; Nguyen, A.; Man, F.Y.; Weinberg, E.; Zeng, Z.; Saltz, L.; Paty, P.B.; Tavazoie, S.F. Extracellular metabolic energetics can promote cancer progression. Cell 2015, 160, 393–406, doi:10.1016/j.cell.2014.12.018.

28. Papalazarou, V.; Zhang, T.; Paul, N.R.; Juin, A.; Cantini, M.; Maddocks, O.D.K.; Salmeron-Sanchez, M.; Machesky, L.M. The creatine-phosphagen system is mechanoresponsive in pancreatic adenocarcinoma and fuels invasion and metastasis. Nat Metab 2020, 2, 62–80, doi:10.1038/s42255-019-0159-z.

29. Kuiper, J.W.; van Horssen, R.; Oerlemans, F.; Peters, W.; van Dommelen, M.M.; te Lindert, M.M.; ten Hagen, T.L.; Janssen, E.; Fransen, J.A.; Wieringa, B. Local ATP generation by brain-type creatine kinase (CK-B) facilitates cell motility. PLoS One 2009, 4, e5030, doi:10.1371/journal.pone.0005030.

30. van Horssen, R.; Janssen, E.; Peters, W.; van de Pasch, L.; Lindert, M.M.; van Dommelen, M.M.; Linssen, P.C.; Hagen, T.L.; Fransen, J.A.; Wieringa, B. Modulation of cell motility by spatial repositioning of enzymatic ATP/ADP exchange capacity. J Biol Chem 2009, 284, 1620–1627, doi:10.1074/jbc.M806974200.

31. Dent, R.; Hanna, W.M.; Trudeau, M.; Rawlinson, E.; Sun, P.; Narod, S.A. Pattern of metastatic spread in triple-negative breast cancer. Breast Cancer Res Treat 2009, 115, 423–428, doi:10.1007/s10549-008-0086-2.

32. Bergnes, G.; Yuan, W.; Khandekar, V.S.; O’Keefe, M.M.; Martin, K.J.; Teicher, B.A.; Kaddurah-Daouk, R. Creatine and phosphocreatine analogs: anticancer activity and enzymatic analysis. Oncol Res 1996, 8, 121–130.

33. Hunter, K.W.; Broman, K.W.; Voyer, T.L.; Lukes, L.; Cozma, D.; Debies, M.T.; Rouse, J.; Welch, D.R. Predisposition to efficient mammary tumor metastatic progression is linked to the breast cancer metastasis suppressor gene Brms1. Cancer Res 2001, 61, 8866–8872.

34. Wang, F.; Samudio, I.; Safe, S. Transcriptional activation of rat creatine kinase B by 17beta-estradiol in MCF-7 cells involves an estrogen responsive element and GC-rich sites. J Cell Biochem 2001, 84, 156–172, doi:10.1002/jcb.1276.

35. Stiehl, D.P.; Bordoli, M.R.; Abreu-Rodriguez, I.; Wollenick, K.; Schraml, P.; Gradin, K.; Poellinger, L.; Kristiansen, G.; Wenger, R.H. Non-canonical HIF-2alpha function drives autonomous breast cancer cell growth via an AREG-EGFR/ErbB4 autocrine loop. Oncogene 2012, 31, 2283–2297, doi:10.1038/onc.2011.417.

36. Balasubramani, M.; Day, B.W.; Schoen, R.E.; Getzenberg, R.H. Altered expression and localization of creatine kinase B, heterogeneous nuclear ribonucleoprotein F, and high mobility group box 1 protein in the nuclear matrix associated with colon cancer. Cancer Res 2006, 66, 763–769, doi:10.1158/0008-5472.CAN-05-3771.

37. Lillie, J.W.; O’Keefe, M.; Valinski, H.; Hamlin, H.A., Jr.; Varban, M.L.; Kaddurah-Daouk, R. Cyclocreatine (1-carboxymethyl-2-iminoimidazolidine) inhibits growth of a broad spectrum of cancer cells derived from solid tumors. Cancer Res 1993, 53, 3172–3178.

38. Martin, K.; Winslow, E.; Okeefe, M.; Khandekar, V.; Hamlin, A.; Lillie, J.; Kaddurahdaouk, R. Specific targeting of tumor cells by the creatine analog cyclocreatine. Int J Oncol 1996, 9, 993–999, doi:10.3892/ijo.9.5.993.

39. Martin, K.J.; Winslow, E.R.; Kaddurah-Daouk, R. Cell cycle studies of cyclocreatine, a new anticancer agent. Cancer Res 1994, 54, 5160–5165.

40. Hoosein, N.M.; Martin, K.J.; Abdul, M.; Logothetis, C.J.; Kaddurah-Daouk, R. Antiproliferative effects of cyclocreatine on human prostatic carcinoma cells. Anticancer Res 1995, 15, 1339–1342.

41. Miller, E.E.; Evans, A.E.; Cohn, M. Inhibition of rate of tumor growth by creatine and cyclocreatine. Proc Natl Acad Sci U S A 1993, 90, 3304–3308, doi:10.1073/pnas.90.8.3304.

42. Fenouille, N.; Bassil, C.F.; Ben-Sahra, I.; Benajiba, L.; Alexe, G.; Ramos, A.; Pikman, Y.; Conway, A.S.; Burgess, M.R.; Li, Q.; et al. The creatine kinase pathway is a metabolic vulnerability in EVI1-positive acute myeloid leukemia. Nat Med 2017, 23, 301–313, doi:10.1038/nm.4283.

43. Kurmi, K.; Hitosugi, S.; Yu, J.; Boakye-Agyeman, F.; Wiese, E.K.; Larson, T.R.; Dai, Q.; Machida, Y.J.; Lou, Z.; Wang, L.; et al. Tyrosine Phosphorylation of Mitochondrial Creatine Kinase 1 Enhances a Druggable Tumor Energy Shuttle Pathway. Cell Metab 2018, 28, 833–847 e838, doi:10.1016/j.cmet.2018.08.008.

44. Tallarida, R.J. An overview of drug combination analysis with isobolograms. J Pharmacol Exp Ther 2006, 319, 1–7, doi:10.1124/jpet.106.104117.

45. Vranic, S.; Gatalica, Z.; Wang, Z.Y. Update on the molecular profile of the MDA-MB-453 cell line as a model for apocrine breast carcinoma studies. Oncol Lett 2011, 2, 1131–1137, doi:10.3892/ol.2011.375.

46. Chevalier, C.; Cannet, A.; Descamps, S.; Sirvent, A.; Simon, V.; Roche, S.; Benistant, C. ABL tyrosine kinase inhibition variable effects on the invasive properties of different triple negative breast cancer cell lines. PLoS One 2015, 10, e0118854, doi:10.1371/journal.pone.0118854.

47. Martin, K.J.; Vassallo, C.D.; Teicher, B.A.; Kaddurah-Daouk, R. Microtubule stabilization and potentiation of taxol activity by the creatine analog cyclocreatine. Anticancer Drugs 1995, 6, 419–426, doi:10.1097/00001813-199506000-00009.

48. Martin, K.H.; Hayes, K.E.; Walk, E.L.; Ammer, A.G.; Markwell, S.M.; Weed, S.A. Quantitative measurement of invadopodia-mediated extracellular matrix proteolysis in single and multicellular contexts. J Vis Exp 2012, e4119, doi:10.3791/4119.

49. de Heer, E.C.; Jalving, M.; Harris, A.L. HIFs, angiogenesis, and metabolism: elusive enemies in breast cancer. J Clin Invest 2020, 130, 5074–5087, doi:10.1172/JCI137552.

50. Gatza, M.L.; Kung, H.N.; Blackwell, K.L.; Dewhirst, M.W.; Marks, J.R.; Chi, J.T. Analysis of tumor environmental response and oncogenic pathway activation identifies distinct basal and luminal features in HER2-related breast tumor subtypes. Breast Cancer Res 2011, 13, R62, doi:10.1186/bcr2899.

51. Network, C.G.A. Comprehensive molecular portraits of human breast tumours. Nature 2012, 490, 61–70, doi:10.1038/nature11412.

52. Lee, J.W.; Bae, S.H.; Jeong, J.W.; Kim, S.H.; Kim, K.W. Hypoxia-inducible factor (HIF-1)alpha: its protein stability and biological functions. Exp Mol Med 2004, 36, 1–12, doi:10.1038/emm.2004.1.

53. Semenza, G.L. Targeting HIF-1 for cancer therapy. Nat Rev Cancer 2003, 3, 721–732, doi:10.1038/nrc1187.

54. Decking, U.K.; Alves, C.; Wallimann, T.; Wyss, M.; Schrader, J. Functional aspects of creatine kinase isoenzymes in endothelial cells. Am J Physiol Cell Physiol 2001, 281, C320–328, doi:10.1152/ajpcell.2001.281.1.C320.

55. Loike, J.D.; Kozler, V.F.; Silverstein, S.C. Creatine kinase expression and creatine phosphate accumulation are developmentally regulated during differentiation of mouse and human monocytes. J Exp Med 1984, 159, 746–757, doi:10.1084/jem.159.3.746.

56. Rahbani, J.F.; Roesler, A.; Hussain, M.F.; Samborska, B.; Dykstra, C.B.; Tsai, L.; Jedrychowski, M.P.; Vergnes, L.; Reue, K.; Spiegelman, B.M.; et al. Creatine kinase B controls futile creatine cycling in thermogenic fat. Nature 2021, 590, 480–485, doi:10.1038/s41586-021-03221-y.

57. Attanasio, F.; Caldieri, G.; Giacchetti, G.; van Horssen, R.; Wieringa, B.; Buccione, R. Novel invadopodia components revealed by differential proteomic analysis. Eur J Cell Biol 2011, 90, 115–127, doi:10.1016/j.ejcb.2010.05.004.

58. Zhang, L.; Zhu, Z.; Yan, H.; Wang, W.; Wu, Z.; Zhang, F.; Zhang, Q.; Shi, G.; Du, J.; Cai, H.; et al. Creatine promotes cancer metastasis through activation of Smad2/3. Cell Metab 2021, 33, 1111–1123 e1114, doi:10.1016/j.cmet.2021.03.009.

59. Chen, Q.; Zhang, N.; Gray, R.S.; Li, H.; Ewald, A.J.; Zahnow, C.A.; Pan, D. A temporal requirement for Hippo signaling in mammary gland differentiation, growth, and tumorigenesis. Genes Dev 2014, 28, 432–437, doi:10.1101/gad.233676.113.

60. Jing, X.; Yang, X.; Zhang, W.; Wang, S.; Cui, X.; Du, T.; Li, T. Mechanical loading induces HIF-1alpha expression in chondrocytes via YAP. Biotechnol Lett 2020, 42, 1645–1654, doi:10.1007/s10529-020-02910-4.

61. Jia, Y.; Li, H.Y.; Wang, J.; Wang, Y.; Zhang, P.; Ma, N.; Mo, S.J. Phosphorylation of 14-3-3zeta links YAP transcriptional activation to hypoxic glycolysis for tumorigenesis. Oncogenesis 2019, 8, 31, doi:10.1038/s41389-019-0143-1.

62. Rabkin, S.W.; Sunga, P. The effect of doxorubicin (adriamycin) on cytoplasmic microtubule system in cardiac cells. J Mol Cell Cardiol 1987, 19, 1073–1083, doi:10.1016/s0022-2828(87)80352-8.

63. Cacciante, F.; Gennaro, M.; Sagona, G.; Mazziotti, R.; Lupori, L.; Cerri, E.; Putignano, E.; Butt, M.; Do, M.T.; McKew, J.C.; et al. Cyclocreatine treatment ameliorates the cognitive, autistic and epileptic phenotype in a mouse model of Creatine Transporter Deficiency. Sci Rep 2020, 10, 18361, doi:10.1038/s41598-020-75436-4.

64. Uemura, T.; Ito, S.; Masuda, T.; Shimbo, H.; Goto, T.; Osaka, H.; Wada, T.; Couraud, P.O.; Ohtsuki, S. Cyclocreatine Transport by SLC6A8, the Creatine Transporter, in HEK293 Cells, a Human Blood-Brain Barrier Model Cell, and CCDSs Patient-Derived Fibroblasts. Pharm Res 2020, 37, 61, doi:10.1007/s11095-020-2779-0.

65. Kurosawa, Y.; Degrauw, T.J.; Lindquist, D.M.; Blanco, V.M.; Pyne-Geithman, G.J.; Daikoku, T.; Chambers, J.B.; Benoit, S.C.; Clark, J.F. Cyclocreatine treatment improves cognition in mice with creatine transporter deficiency. J Clin Invest 2012, 122, 2837–2846, doi:10.1172/JCI59373.

66. Chang, E.J.; Ha, J.; Oerlemans, F.; Lee, Y.J.; Lee, S.W.; Ryu, J.; Kim, H.J.; Lee, Y.; Kim, H.M.; Choi, J.Y.; et al. Brain-type creatine kinase has a crucial role in osteoclast-mediated bone resorption. Nat Med 2008, 14, 966–972, doi:10.1038/nm.1860.

67. (NCATS), N.N.C.f.A.T.S. LUM-001 as a treatment for creatine transporter deficiency. Available online: https://ncats.nih.gov/trnd/projects/complete/cincy-creatine-transporter-defect (accessed on August 19, 2021).

68. Brooks, D.L.; Seagroves, T.N. Chromatin Immunoprecipitation of HIF-alpha in Breast Tumor Cells Using Wild Type and Loss of Function Models. Methods Mol Biol 2018, 1742, 67–79, doi:10.1007/978-1-4939-7665-2_7.

69. Seagroves, T.N.; Ryan, H.E.; Lu, H.; Wouters, B.G.; Knapp, M.; Thibault, P.; Laderoute, K.; Johnson, R.S. Transcription factor HIF-1 is a necessary mediator of the pasteur effect in mammalian cells. Mol Cell Biol 2001, 21, 3436–3444, doi:10.1128/MCB.21.10.3436-3444.2001.

70. Deng, S.; Krutilina, R.I.; Wang, Q.; Lin, Z.; Parke, D.N.; Playa, H.C.; Chen, H.; Miller, D.D.; Seagroves, T.N.; Li, W. An Orally Available Tubulin Inhibitor, VERU-111, Suppresses Triple-Negative Breast Cancer Tumor Growth and Metastasis and Bypasses Taxane Resistance. Mol Cancer Ther 2020, 19, 348–363, doi:10.1158/1535-7163.MCT-19-0536.

## Supplementary Materials and Methods References

1. Schwab, L.P.; Peacock, D.L.; Majumdar, D.; Ingels, J.F.; Jensen, L.C.; Smith, K.D.; Cushing, R.C.; Seagroves, T.N. Hypoxia-inducible factor 1alpha promotes primary tumor growth and tumor-initiating cell activity in breast cancer. Breast Cancer Res 2012, 14, R6, doi:10.1186/bcr3087.

2. Network, C.G.A. Comprehensive molecular portraits of human breast tumours. Nature 2012, 490, 61–70, doi:10.1038/nature11412.

3. Nagy, A.; Lanczky, A.; Menyhart, O.; Gyorffy, B. Validation of miRNA prognostic power in hepatocellular carcinoma using expression data of independent datasets. Sci Rep 2018, 8, 9227, doi:10.1038/s41598-018-27521-y.

4. Brooks, D.L.; Schwab, L.P.; Krutilina, R.; Parke, D.N.; Sethuraman, A.; Hoogewijs, D.; Schorg, A.; Gotwald, L.; Fan, M.; Wenger, R.H.; et al. ITGA6 is directly regulated by hypoxia-inducible factors and enriches for cancer stem cell activity and invasion in metastatic breast cancer models. Mol Cancer 2016, 15, 26, doi:10.1186/s12943-016-0510-x.

5. Oosthuyse, B.; Moons, L.; Storkebaum, E.; Beck, H.; Nuyens, D.; Brusselmans, K.; Van Dorpe, J.; Hellings, P.; Gorselink, M.; Heymans, S.; et al. Deletion of the hypoxia-response element in the vascular endothelial growth factor promoter causes motor neuron degeneration. Nature genetics 2001, 28, 131–138, doi:10.1038/88842.

6. Choi, B.H.; Ha, Y.; Ahn, C.H.; Huang, X.; Kim, J.M.; Park, S.R.; Park, H.; Park, H.C.; Kim, S.W.; Lee, M. A hypoxia-inducible gene expression system using erythropoietin 3′ untranslated region for the gene therapy of rat spinal cord injury. Neurosci Lett 2007, 412, 118–122, doi:10.1016/j.neulet.2006.11.015.

7. Martin, K.H.; Hayes, K.E.; Walk, E.L.; Ammer, A.G.; Markwell, S.M.; Weed, S.A. Quantitative measurement of invadopodia-mediated extracellular matrix proteolysis in single and multicellular contexts. J Vis Exp 2012, e4119, doi:10.3791/4119.

8. Tallarida, R.J. An overview of drug combination analysis with isobolograms. J Pharmacol Exp Ther 2006, 319, 1–7, doi:10.1124/jpet.106.104117.

