## Supplementary Tables S4-S7 for "HIF-dependent CKB expression promotes breast cancer metastasis, whereas cyclocreatine therapy impairs cellular invasion and improves chemotherapy efficacy"

**Table S4**. Primers and Roche Universal Probe Library (UPL) FAM-labeled probes utilized in real-time PCR assays. All assays were designed using the Roche Universal ProbeLibrary Assay Design Center.

| Gene | Forward Primer | Reverse Primer | UPL ID |
|---|---|---|---|
| *Ckb (murine)* | gcaagcacaggcatccat | cgcagcttctgcgtattatg | 70 |
| *CKB (human)* | cctgcccagaaatgaagc | gcactgcccaggcaataa | 38 |
| *Ints3 (murine)* | gtggctgttattgactctgcac | caggttccccatcatcacat | 17 |
| *Krt18* (murine K18) | agatgacaccaacatcacaagg | cttccagaccttggacttcct | 78 |
| *PPIA (Cyclophilin A)* | atgctggacccaacacaaat | tctttcactttgccaaacacc | 48 |

**Table S5**. Primary antibody source and dilution factors utilized in western blotting, immunofluorescence (IF) staining of cultured cells and DAB-based immunohistochemistry (IHC) staining of tumor tissues.

| Antibody | Source and Catalog # | Dilution | Purpose |
|---|---|---|---|
| anti-CKB | AbCAM (#ab88746) | 1:5,000 | western |
| anti-ß-tubulin | AbCam (#(#ab6046) | 1:10,000 | western |
| anti-HIF-1$\alpha$ | Novus Biologicals (#NB100-479) | 1:5,000 | western |
| anti-PARP | Cell Signaling Technology (#9542) | 1:5,000 | western |
| anti-CKB | ThermoFisher (PA5-21382) or Santa Cruz, sc-271531) | 1:500 | IF, cells |
| anti-alpha tubulin | ThermoFisher (#62204) | 1:250 | IF, cells |
| anti-cortactin | Millipore (#05-0180) | 1:700 | IF, cells |
| anti-mouse AlexaFluor-488 | LifeTech (#A21202) | 1:400 | IF, cells |
| anti-rabbit AlexaFluor-594 | LifeTech (#A21203) | 1:400 | IF, cells |
| anti-rabbit-IgG-Biotin-X | LifeTech (#A16027) | 1:400 | IF, cells |
| Strepavidin-AlexaFlour-594 | LifeTech (#S11227) | 1:400 | IF, cells |
| anti-Ki67 | AbCAM (#ab-15580) | 1:750 | IHC |
| anti-CD31 | Cell Signaling Technology (#77699) | 1:100 | IHC |
| anti-activated caspase3 | Cell Signaling Technology (#9661) | 1:100 | IHC |

**Table S6**. Primers used in chromatin immunoprecipitation (ChIP) assays in murine PyMT tumor cells and in human MCF-7 cells.

| Genomic Region | IP Antibody | Forward Primer | Reverse Primer |
|---|---|---|---|
| -258 *CKB* | HIF-1α | gatgaacccaagcgtcctc | agacctcgaggccgaaac |
| -935 *CKB* | HIF-1α | attgctgggtcacggagtt | tacccccaacactcccagat |
| -1869 *CKB* non-HRE | HIF-1α | ttcaggttttgtggggtagc | ctgttccaagccagcatttt |
| -1326 *Ckb* | HIF-1α | cagggctgttctggactctc | cctcacaagtgctgggatta |
| -1835 *Ckb* | HIF-1α | tgcctcctgtgcacttttt | agcctagcttcagagaactaa tgg |
| -510 *Ckb* non-HRE | HIF-1α | cttggcggtgttccttagag | caggccatatcctcaagagc |
| *EPO* | HIF-1α | gctggcctctggctctcatgg | cagggttggcagctgccttact g |
| *Vegf* | HIF-1α | ctggcttcagttccctggcaacat ctct | cctggggtgaatgggatcctct gg |

**Table S7**. *Ckb* shRNA targeting sequences obtained from The RNAi Consortium (TRC) library purchased through Open Biosystems.

| Clone ID | Clone Name | Target Sequence |
|---|---|---|
| TRCN0000024659 | NM_021273.2-300s1c1 | cgacgtattcaaggacctctt |
| TRCN0000024660 | NM_021273.2-288s1c1 | cgaggagagttacgacgtatt |
| TRCN0000024661 | NM_021273.2-708s1c1 | gcacaatgacaataagacttt |
