## Supplementary material for "HIF-dependent CKB expression promotes breast cancer metastasis, whereas cyclocreatine therapy impairs cellular invasion and improves chemotherapy efficacy": SuppFigures and Legends

FIG. S1

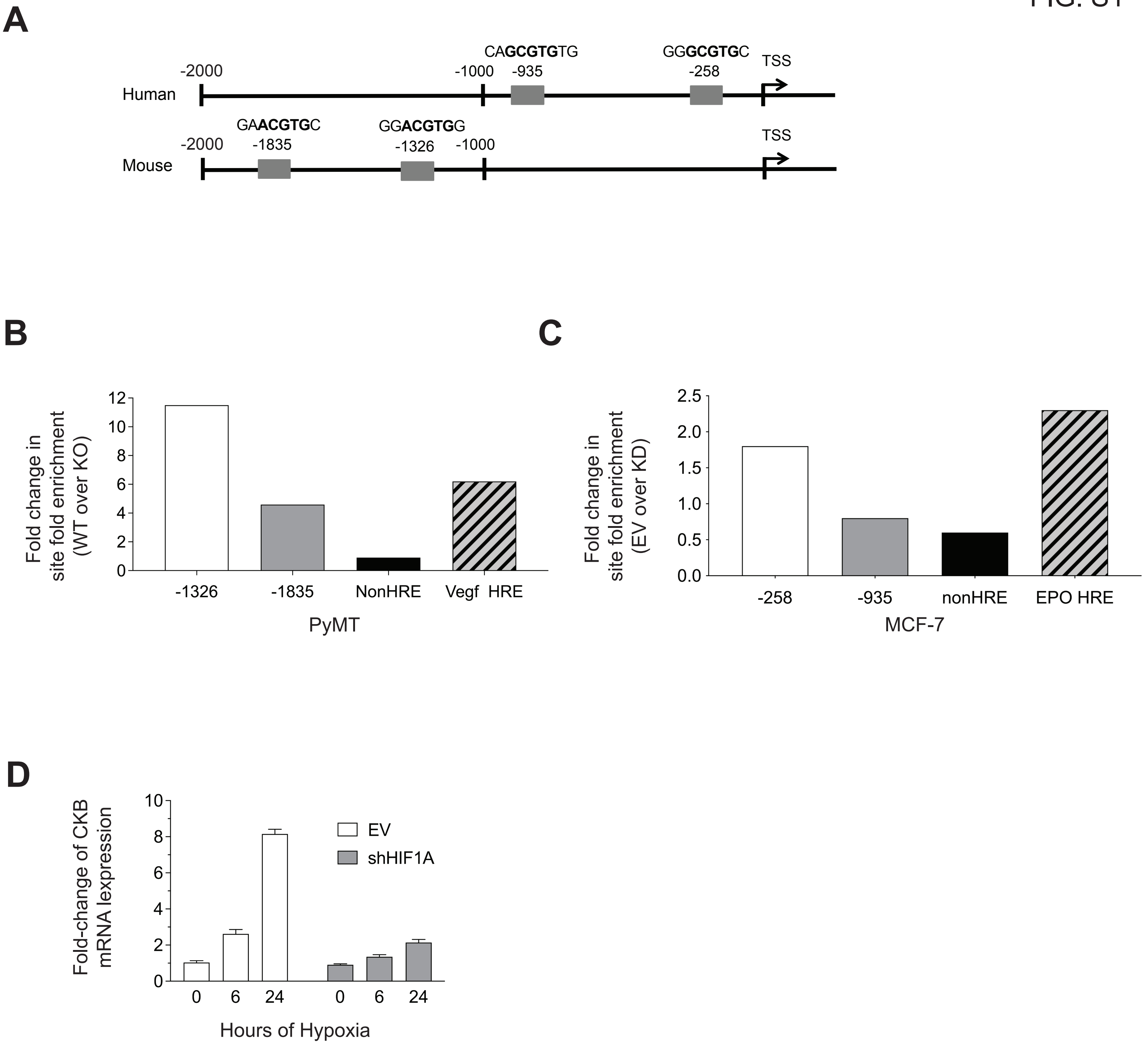
A
CAGCGTGTG
GGGCGTGC
-2000
-1000
-935
-258
TSS
Human
GAACGTGC
GGACGTGG
-2000
-1835
-1326
-1000
TSS
Mouse
B
Fold change in site fold enrichment (WT over KO)
12
10
8
6
4
2
0
-1326
-1835
NonHRE
Vegf HRE
PyMT
C
Fold change in site fold enrichment (EV over KD)
2.5
2.0
1.5
1.0
0.5
0.0
-258
-935
nonHRE
EPO HRE
MCF-7
D
Fold-change of CKB mRNA lexpression
10
8
6
4
2
0
EV
shHIF1A
0
6
24
0
6
24
Hours of Hypoxia

A

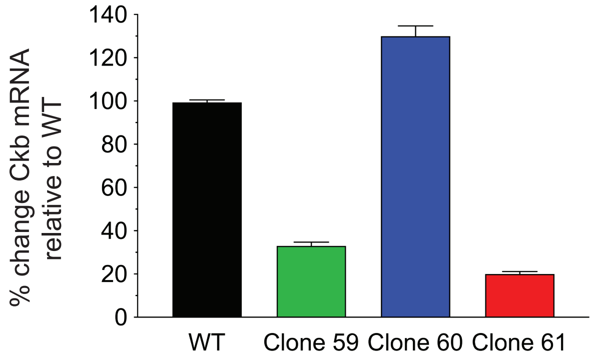
140
120
100
80
60
40
20
0
% change Ckb mRNA relative to WT
WT
Clone 59
Clone 60
Clone 61

B

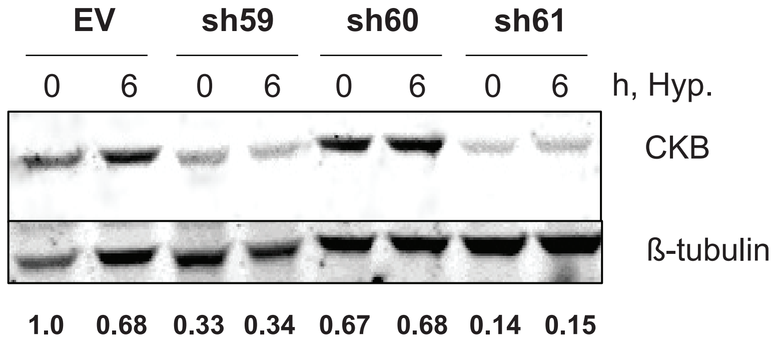
EV
sh59
sh60
sh61
0 6 0 6 0 6 0 6 h, Hyp.
CKB
ß-tubulin
1.0 0.68 0.33 0.34 0.67 0.68 0.14 0.15

C

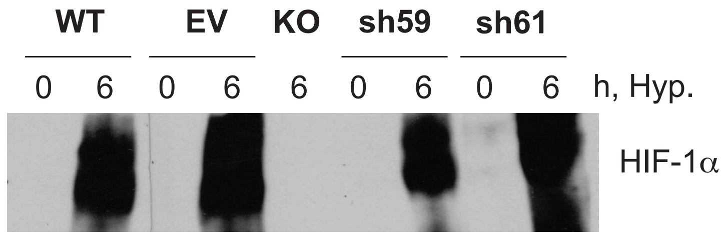
WT
EV
KO
sh59
sh61
0 6 0 6 6 0 6 0 6 h, Hyp.
HIF-1α

FIG. S3

A

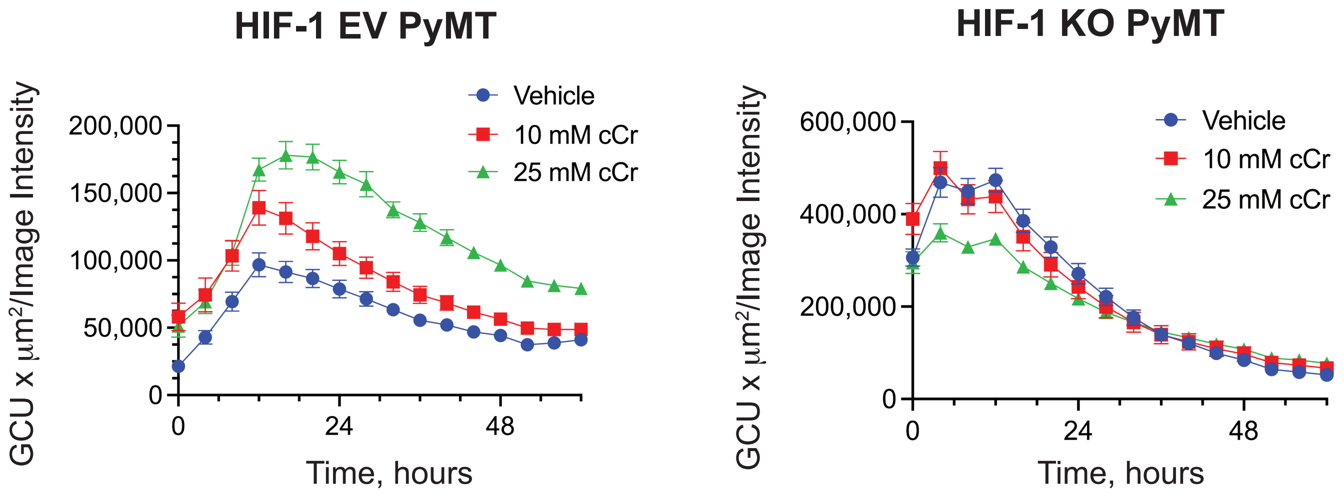

HIF-1 EV PyMT
GCU x μm²/Image Intensity
200,000
150,000
100,000
50,000
0
0
24
48
Time, hours
Vehicle
10 mM cCr
25 mM cCr
HIF-1 KO PyMT
GCU x μm²/Image Intensity
600,000
400,000
200,000
0
0
24
48
Time, hours
Vehicle
10 mM cCr
25 mM cCr


B

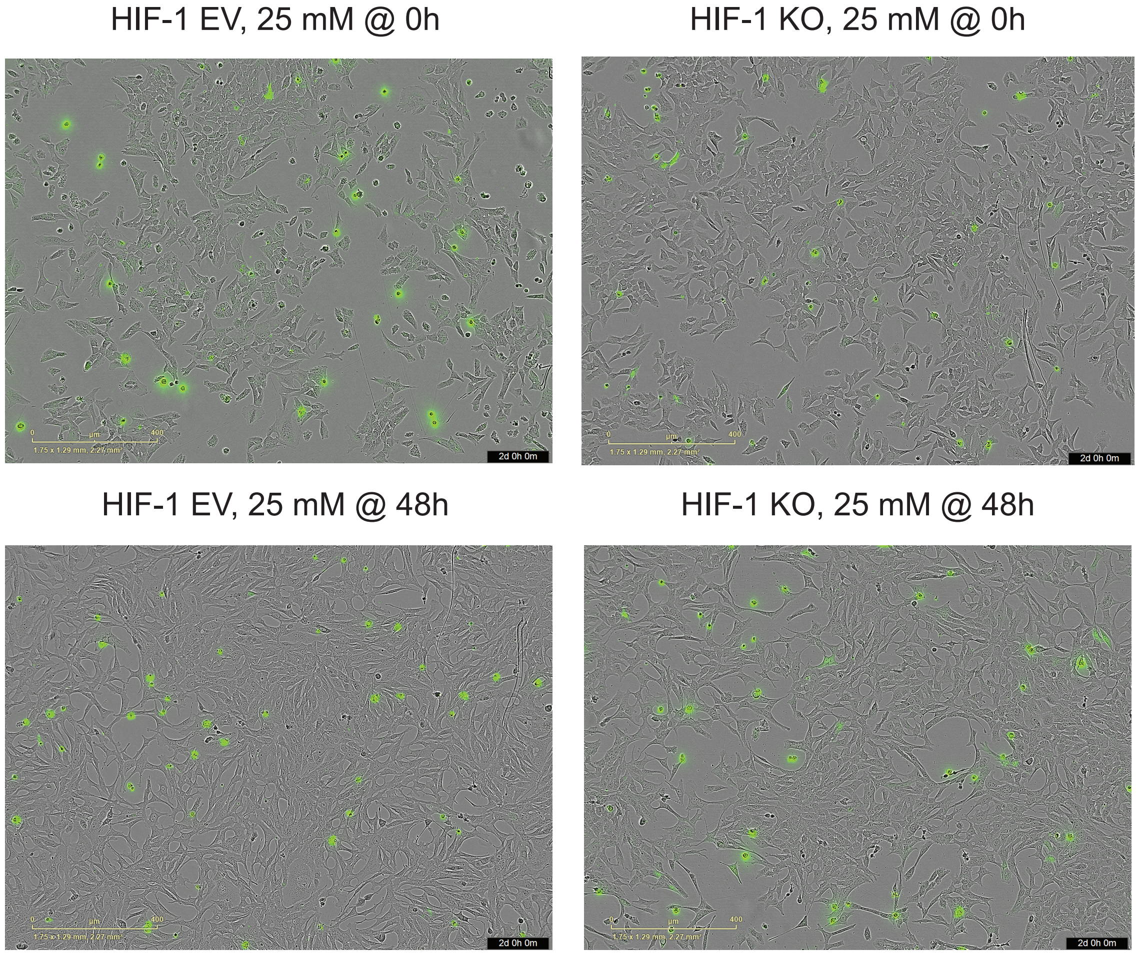

HIF-1 EV, 25 mM @ 0h
HIF-1 KO, 25 mM @ 0h
HIF-1 EV, 25 mM @ 48h
HIF-1 KO, 25 mM @ 48h

FIGURE S4

**A**

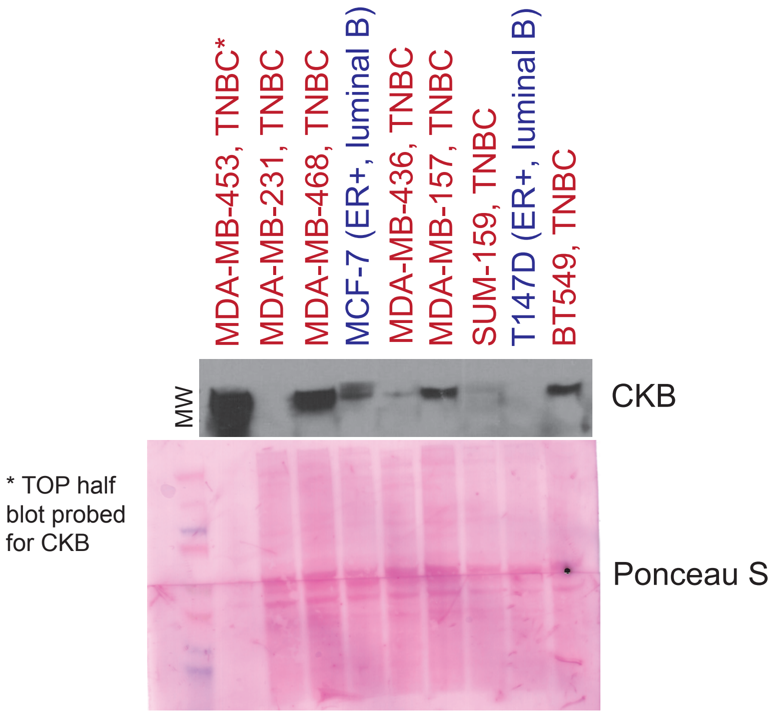


*MDA-MB-453 cells underloaded by 5-fold due to high levels of expression

FIGURE S5

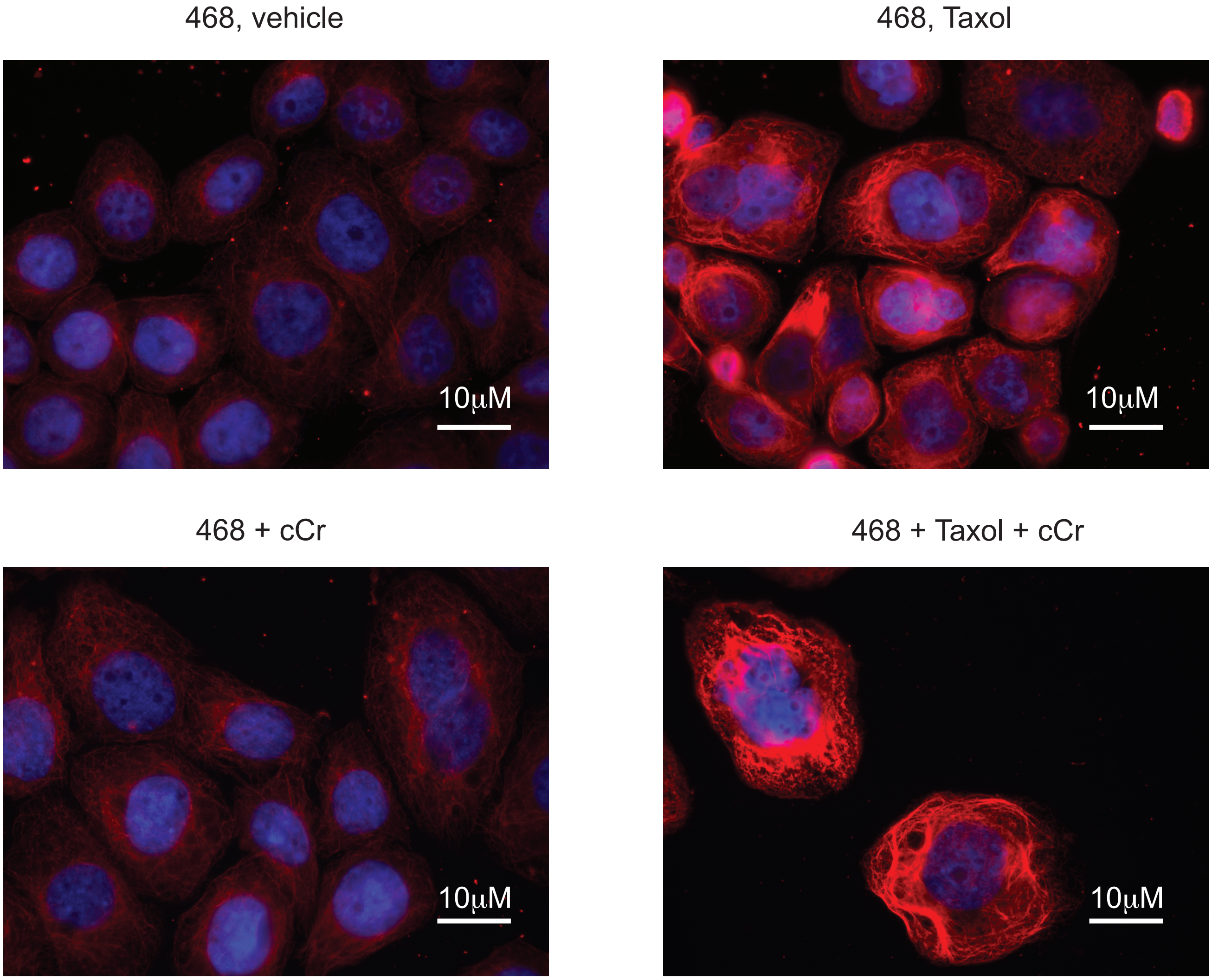
468, vehicle
10μM
468, Taxol
10μM
468 + cCr
10μM
468 + Taxol + cCr
10μM

FIG. S6

**A**

BT549, vehicle BT549 + cCr

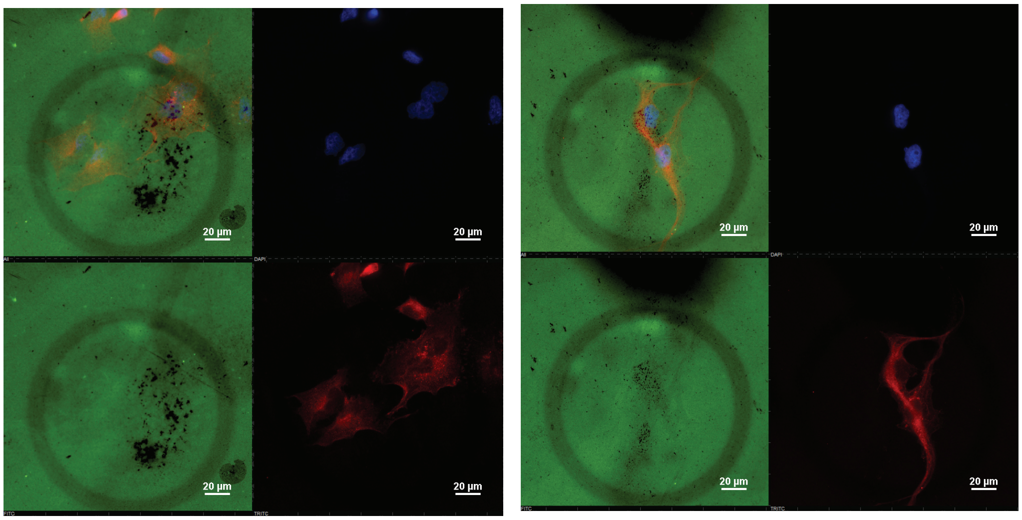


gelatin
cortactin
DAPI

**B**

231 cells +mCKB

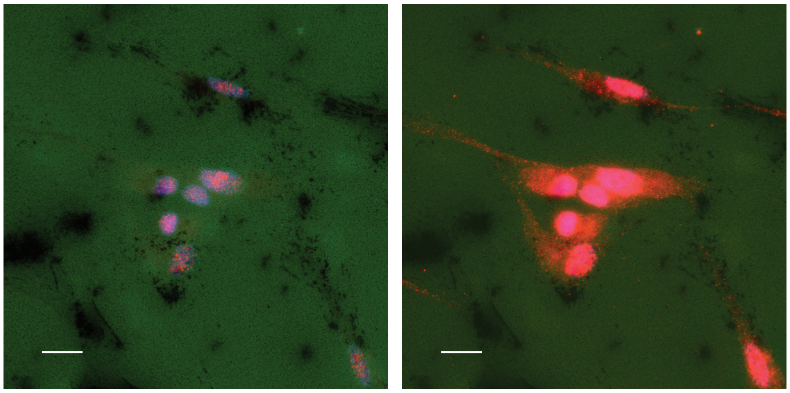

gelatin
mCKB
DAPI

short exposure long exposure

**Supplementary Figure Legends**

**Supplementary Figure S1. Chromatin immunoprecipitation (ChIP) analysis of the *Ckb/CKB* proximal promoter in PyMT and MCF-7 breast cancer cells.**

A. A schematic representation of two consensus putative hypoxia response elements (HREs) identified by the JASPAR database in either the mouse (*Ckb*) or the human (*CKB*) gene proximal promoter. B-C. All cells were cultured to ~80% confluence and then exposed to hypoxia for 6-24h (0.5% $O_2$) prior to the isolation of chromatin. ChIP assays were performed with antibodies to HIF-1$\alpha$ (AbCAM, cat# H1alpha67, ab1) and qPCR was conducted on the isolated DNA to determine HIF1$\alpha$ recruitment to HREs. Binding enrichment represented at each HRE site is expressed as the fold change between HIF-1 WT PyMT cells (B), or MCF-7 empty vector (EV) (C) cells relative to either HIF-1 KO PyMT cells (B) or MCF-7 shHIF1A knockdown (KD) cells (C), respectively, after normalizing to the IgG antibody control. As internal assay controls for the PyMT cells, ChIP was also performed for a known HRE in the mouse *Vegf* promoter [5] or using an intron within *Ckb* with no identified HRE sites. As internal assay controls for MCF-7 cells, ChIP for HIF-1$\alpha$ was performed in a known HRE site in the *EPO* 3' UTR [6], or using an intron within *CKB* with no identified HRE sites. D. *CKB* mRNA levels were measured by qPCR in MCF-7 EV and shHIF1A cells cultured for 0, 6 or 24h of hypoxia. The fold change in *CKB* expression was calculated after normalization for loading to *PPIA* (n=3 technical replicates/cell line/condition; data are representative of two independent experiments).

**Supplementary Figure S2. Comparing gene knockdown efficiency of *Ckb* shRNA constructs in PyMT cells.**

A. HIF-1 WT PyMT cells were transfected with three different pLKO.1-puro-shRNA targeting viral backbones to screen for deletion efficiency at the mRNA level by qPCR (n=4 technical replicates/cell line) when cells were cultured at normoxia. B. Western blot analysis of CKB protein levels using replicate plates from (A), along with EV (empty vector) PyMT cells (WT cells transduced to express empty pLKO.1-puro); ß-tubulin is used as a loading control. The EV 0h hypoxia sample was set to 1.0 and ImageJ was used to evaluate densitometry values of CKB protein. C. Western blotting for HIF-1$\alpha$ in WT parental, HIF-1 WT + EV (EV), HIF-1 KO, or *Ckb* sh59 KD and sh61 KD pools cultured at normoxia (excluding HIF-1 KO) or for 6h at hypoxia following transduction with pLKO.1-puro-based recombinant lentiviruses and the selection of stable shRNA KD pools using puromycin. Western blots for HIF-1$\alpha$ were performed as in [1]. HIF-1$\alpha$ is present as a poly-ubiquitinylated protein smear after exposure to hypoxia; the 6h HIF-1 KO sample serves as an internal negative control.

**Supplementary Figure S3. CytoTox Green assays to measure cell viability after exposure to 15 mM or 25 mM cCr.**

A. PyMT EV cells used in wound healing, invasion, and cell cycle progression experiments as presented in Figure 4, or HIF-1 KO cells, were grown in 2% FBS (standard growth medium) and then exposed to either 15 mM cCr (the ~$IC_{50}$ dose @96h of exposure) or 25 mM cCr in the presence of CytoTox Green reagent. The intensity of CytoTox Green dye incorporated by dead cells was enumerated over time using the IncuCyte S3 live-cell imager. HIF-1 KO cells have a higher basal level of cell death relative to EV cells on a per area basis at t=0 h. B. Example well scan images for each genotype are shown at t=0h (1 h after addition of CytoTox and when scanning began) and at t=48h (experimental endpoint). Overall, these images indicate that the frequency of CytoTox Green positive cells is approximately equivalent for PyMT EV and PyMT HIF-1 KO genotypes after 48h of treatment with 25 mM cCr, relative to their respective t=0h timepoints, and that the majority of cells remain viable after 48h of treatment.

**Supplementary Figure S4. Comparison of CKB levels in additional human breast cancer cell lines.** A. Western blot comparing CKB expression between TNBC cells (red font: MDA-MB-453, MDA-MB-231, MDA-MB-468, MDA-MB-436, MDA-MB-157, SUM-159, BT-549) and two ER-positive luminal B models (blue font), MCF-7 cells, known to express CKB, and T47D cells. Of note, the MDA-MB-453 sample was underloaded by ~5-fold due to very high levels of expression (refer to Figure 7A). For the other samples, loading is approximately equivalent as shown by Ponceau S staining of the whole blot; however, only the top half of the blot was probed for CKB.

**Supplementary Figure S5. Tubulin networks detected by alpha-tubulin immunostaining following paclitaxel (Taxol) treatment in combination with cCr.**

MDA-MB-468 cells were plated into ibidi multi-chamber well #1.5H glass slides, allowed to adhere for 48h, treated with either 20 mM cCr, 10 nM Taxol, or both drugs for 48h, then washed, fixed, and immunostained with alpha-tubulin (red) and counterstained with DAPI. All images were captured on a Nikon ECLIPSE Ti2 microscope at the same laser intensities and exposure times at 400x magnification; scale bars represent 10 $\mu$M.

**Supplementary Figure S6. Immunostaining of invadopodia coverslips at experimental endpoint.**

A. Example images taken from coverslips used for invadopodia assays for the BT549 TNBC cell line in Figure 7. During invadopodia formation, BT549 cells were exposed to either vehicle (left panel) or to 25 mM cCr (~$IC_{50}$; right panel) for 48-50h (green=gelatin; red=cortactin; blue=DAPI). Scale bar represents 20 $\mu$M. B. MDA-MB-231 cells ectopically expressing mCKB readily formed invadopodia (refer to Figure 7). At the study endpoint, coverslips containing invadopodia were immunostained with CKB antibodies. The image represents the same area imaged with either a short or a long exposure time in the red channel with the green laser intensity also minimized. In the longer exposure image, mCKB can be found along the periphery of the cell and in the cell projections overlapping with invadopodia, as well as in the cytoplasm and in the nucleus; the scale bar represents 20 $\mu$M.
